# Anillin forms linear structures and facilitates furrow ingression after septin and formin depletion

**DOI:** 10.1101/2022.12.22.521621

**Authors:** Mikhail Lebedev, Fung-Yi Chan, Jennifer Bellessem, Daniel S. Osório, Elisabeth Rackles, Tamara Mikeladze-Dvali, Ana X. Carvalho, Esther Zanin

**Author notes:** Correspondence to: Esther Zanin.

## Abstract

During cytokinesis a contractile ring consisting of unbranched filamentous actin (F-actin) and myosin II filaments assembles and constricts at the cell equator. Unbranched F-actin is *de novo* generated by formin and without formin cleavage furrow ingression fails. In *C. elegans* depletion of septin restores cleavage furrow ingression in formin (CYK-1) mutants. How the cleavage furrow ingresses without a detectable unbranched F-actin ring is not known. We report, that in this setting anillin (ANI-1) is essential for furrow ingression and forms a meshwork of linear structures, which circumferentially align around the cell equator. Although equatorial ANI-1 recruitment is facilitated by septins, the formation of linear ANI-1 structures is septin independent. Analysis of ANI-1 deletion mutants reveals that its disordered linker region is required for linear structure formation and furrow ingression. We also found that myosin II (NMY-2) decorates linear ANI-1 structures and promotes their circumferential alignment. NMY-2 also interacts with various lipids and forms membrane localized clusters in absence of F-actin and anillin binding. This suggests that NMY-2 represents an independent link between the F-actin / ANI-1 network and the plasma membrane. Collectively, our data reveals a compensatory mechanism, mediated by ANI-1 linear structures and membrane-bound NMY-2, that promotes furrow formation and ingression when formins are depleted and therefore unbranched F-actin polymerization is compromised.

## Introduction

During cytokinesis a contractile ring consisting of unbranched filamentous actin (F-actin) and myosin II assembles underneath the plasma membrane between the segregating chromosomes. Constriction of the actin-myosin ring splits the mother cell into two daughter cells (D’Avino et al., 2015; Mishima, 2016; Pollard and O’Shaughnessy, 2019). Stimulatory and inhibitory signals emanating from the mitotic spindle activate RhoA at the cell equator, which in turn induces contractile ring assembly (Basant and Glotzer, 2018; Dechant and Glotzer, 2003; Gomez-Cavazos et al., 2020; Mangal et al., 2018; Prokopenko et al., 1999; Schneid et al., 2021; Yüce et al., 2005; Zanin et al., 2013). Active RhoA releases the autoinhibitory conformation of formin with the help of the contractile ring component anillin and IQGAP1 (Chen et al., 2020; Chen et al., 2017; Li and Higgs, 2003; Otomo et al., 2005). Active formin binds to the barbed end of actin filaments and induces the polymerization of long unbranched F-actin (Kühn and Geyer, 2014). F-actin circumferentially aligns around the cell equator which is thought to facilitate furrow ingression (Henson et al., 2017; Leite et al., 2020; Reymann et al., 2016; Schroeder, 1968; Spira et al., 2017).Without formin cleavage furrow formation and cytokinesis fail (Pelham and Chang, 2002; Severson et al., 2002; Swan et al., 1998; Watanabe et al., 2008).

In addition to formin, RhoA also recruits and activates non-muscle myosin II (NMY-2 in *C. elegans*) via RhoA kinase. The motor myosin II interacts with unbranched F-actin and is required for cytokinesis in many systems (Cuenca et al., 2003; De Lozanne and Spudich, 1987; Straight et al., 2003). Myosin II consists of a globular motor domain and coiled-coil domain through which it forms bipolar minifilaments. To be active, myosin II has to bind the phosphorylated regulatory light chain (RLC) and essential light chain (ELC) (Pollard, 2020). In addition to RhoA-dependent myosin II recruitment, a positive feedback mechanism between myosin II generated equatorial compression and equatorial directed flow enhances myosin II accumulation at the cell equator further (Khaliullin et al., 2018). Myosin II becomes enriched at the cell equator independently of its binding partner F-actin or anillin (Hickson and O’Farrell, 2008; Maddox et al., 2005). Myosin II inhibition does not prevent the recruitment of other ring components to the equator but blocks ring constriction (Davies et al., 2014; Straight *et al*., 2003; Tse et al., 2011). In fission yeast myosin II is targeted to the membrane by the anillin homologue Mid1p and together with formin and numerous other proteins they form defined protein assemblies called nodes. Within each node myosin II heads point away from the membrane and grab and pull on actin filaments generated and anchored at adjacent nodes. This leads to the coalescence of nodes and eventually ring constriction (Pollard and O’Shaughnessy, 2019).

Another component of the contractile ring is the conserved multidomain protein anillin (ANI-1 in *C. elegans)* (D’Avino *et al*., 2015; Mishima, 2016; Naydenov et al., 2020; Piekny and Maddox, 2010). Due to its binding to the membrane or membrane-associated proteins, F-actin and myosin II, anillin represents an important linker between the actin-myosin ring and the membrane. Anillin is also considered a scaffolding component of the contractile ring since numerous interaction partners have been documented. With its C-terminus anillin interacts with microtubules (van Oostende Triplet et al., 2014), with lipids through the C2 domain, with membrane-associated active RhoA via the RhoA binding domain (RBD) (Piekny and Glotzer, 2008; Sun et al., 2015), and with septins via the PH domain (El Amine et al., 2013; Field et al., 2005; Liu et al., 2012; Oegema et al., 2000). With its N-terminus anillin binds to myosin II (Straight et al., 2005), F-actin (Field and Alberts, 1995; Jananji et al., 2017; Matsuda et al., 2019; Oegema *et al*., 2000; Tian et al., 2015), formins (Chen *et al*., 2017; Watanabe *et al*., 2008) and importin (Beaudet et al., 2020). Anillin localization to the cell equator requires RhoA activation but not myosin II, F-actin, septins or formins (D’Avino et al., 2008; Hickson and O’Farrell, 2008; Maddox *et al*., 2005; Straight *et al*., 2005; Watanabe *et al*., 2008). Consistent with its role as a membrane-actomyosin linker, anillin depletion results in unstable furrows and in cytokinesis failure in fly and human cells (Piekny and Glotzer, 2008; Straight *et al*., 2005). *C. elegans* has three anillin homologues (ANI-1, ANI-2, and ANI-3) of which only ANI-1 is expressed in the early embryo and required for embryonic development. ANI-1 depletion in zygotes does not cause cytokinesis failure but results in symmetric furrow ingression (Maddox *et al*., 2005; Maddox et al., 2007). Anillin positively feeds back on RhoA activity (Piekny and Glotzer, 2008) by stabilizing active RhoA at the membrane and thereby facilitating RhoA effector binding (Budnar et al., 2019). Anillin also binds and recruits septins to the contractile ring (Field *et al*., 2005; Hickson and O’Farrell, 2008; Liu *et al*., 2012; Maddox *et al*., 2005). Recent *in vitro* work suggests that anillin could contribute to force generation during constriction by passive diffusion along cross linked F-actin filaments (Kučera et al., 2021).

Besides F-actin and myosin II, septins are the third filament forming protein of the contractile ring and septins bind the plasma membrane (Mostowy and Cossart, 2012). Similar to anillin, septins are required for ring ingression in human cells (Estey et al., 2010) but are not essential in *C. elegans* zygotes. Whereas humans have 13 septins, *C. elegans* has two (UNC-59 and UNC-61), which are only essential for some postembryonic divisions (Nguyen et al., 2000). Together, septin and anillin also function during the late stages of cytokinesis during intercellular bridge formation (Estey *et al*., 2010; Green et al., 2013; Kechad et al., 2012; Renshaw et al., 2014).

During the initial assembly phase ring components such as F-actin, myosin II and anillin localize in a broad equatorial band, which subsequently compacts and matures into the contractile ring (Lewellyn et al., 2010). Recent super-resolution microscopy in sea urchin embryos revealed that myosin II, anillin and septin initially co-localize in puncta reminiscent to the nodes in yeast (Chelsea Garno, 2021). Subsequently, a dense meshwork forms in which myosin II filaments orient parallel to actin filaments (Beach et al., 2014; Fenix et al., 2016; Henson *et al*., 2017). This organization is consistent with myosin II mediating ring constriction by actin filament sliding in a purse-string-like fashion. Consistently myosin II motor activity is required for ring constriction (Osório et al., 2019).

Although actin-based contractility appears to be the predominant mechanisms for furrow ingression, alternative mechanisms were also reported sporadically. For example, the migration of the two daughter cells away from each other supports furrow ingression in the absence of a visible F-actin ring in non-transformed human RPE1 cells and in *Dictyostelium* (Dix et al., 2018; Nagasaki et al., 2009; Neujahr et al., 1997) and in *Chlamodymonas* furrow ingression does not require F-actin (Onishi et al., 2020). In *C. elegans* depletion of septin in the formin (CYK-1) temperature sensitive (ts) mutant restored furrow ingression although no F-actin ring was detectable (Jordan et al., 2016). Here we investigate the mechanism of furrow ingression after co-depletion of CYK-1 and septins in the *C. elegans* one-cell embryo. We discover that ANI-1 forms a meshwork of linear structures that circumferentially align around the cell equator and promote furrow ingression without detectable unbranched F-actin and septins. Mutational analysis reveals that the disordered N-terminal half of ANI-1 is sufficient for linear structure formation and furrow ingression. Thereby our work uncovers a novel ANI-1-dependent mechanism of cleavage furrow ingression.

## Results

### ANI-1 is required for furrow ingression after CYK-1 and septin co-depletion

*C. elegans* has six formins and CYK-1, a Diaphanous-type formin, is the only one known to be essential for cytokinesis (Pruyne, 2016; Severson *et al*., 2002; Swan *et al*., 1998). Surprisingly, it was found that *cyk-1ts* mutant embryos, which lack unbranched F-actin, still ingress a cleavage furrow when septins are depleted (Fig. 1A) (Davies et al., 2018; Jordan *et al*., 2016). We confirmed this observation as CYK-1 depletion by RNAi prevented cleavage furrow formation but septin^UNC-61^ co-depletion facilitated cleavage furrow ingression in all embryos (Fig. 1B, C, Movie 1). In 5/10 *cyk-1(RNAi);septin^unc-61^(RNAi)* embryos the furrow remained closed until the onset of the second cell division. In the remaining embryos, the furrow eventually regressed around 800 sec after NEBD (Fig. 1B). *C. elegans* has two septins (UNC-61 and UNC-59) and they depend on each other for membrane localization (Nguyen et al., 2000), therefore depletion of one is sufficient to disrupt the function of the complex. Consistently, we observed that endogenously GFP-tagged septin^UNC-59^ did not localize to the equatorial cortex after *septin^unc-61^ (RNAi)* or *cyk-1(RNAi);septin^unc-61^ (RNAi)* (Fig. S1A, B). Similarly, endogenously tagged CYK-1 or F-actin, visualized by LifeAct::RFP, no longer accumulated at the cell equator after single or double *cyk-1(RNAi)* treatments, confirming previous observations (Chan et al., 2018; Davies *et al*., 2018; Ding et al., 2017; Jordan *et al*., 2016) (Fig. S1C, D). Immunoblot analysis of adult worms revealed that *septin^unc-59^(RNAi)* and *cyk-1(RNAi)* strongly reduced endogenously-tagged septin^UNC-59^ and CYK-1 protein levels, respectively (Fig. S1E). In addition to formin nucleated F-actin, the cortex contains Arp2/3 nucleated branched F-actin, which is not essential for cytokinesis (Chan *et al*., 2018). Thus, we tested whether the remaining Arp2/3-generated F-actin contributes to furrowing in CYK-1 and septin co-depleted embryos by additional depletion of the Arp2 homologue ARX-2. All *cyk-1(RNAi); septin^unc-61^(RNAi); arx-2(RNAi)* embryos fully ingressed a cleavage furrow (Fig. S2A, B), suggesting that Arp2/3-generated F-actin is also not essential for furrowing in this condition. To address the mechanism how CYK-1 and septin co-depleted embryos ingress a cleavage furrow, we determined whether ANI-1 is required for furrow ingression in this background. Whereas ANI-1 depletion alone does not cause cytokinesis failure in the one-cell embryo (Maddox *et al*., 2005), ANI-1 depletion together with CYK-1 and septin^UNC-61^ prevented cleavage furrow ingression (Fig. 1B, C). Immunoblot analysis of ANI-1 protein levels showed that ANI-1 levels were strongly reduced (Fig. S2C). Further, quantification of ANI-1 fluorescence intensity on the cortex in *ani-1* single, double and triple RNAi depletions confirmed that ANI-1 depletion was highly efficient in the one-cell embryo (Fig. S2D, E). Together, ANI-1 promotes furrow ingression in case CYK-1 and septin levels are low.

**Figure 1.**
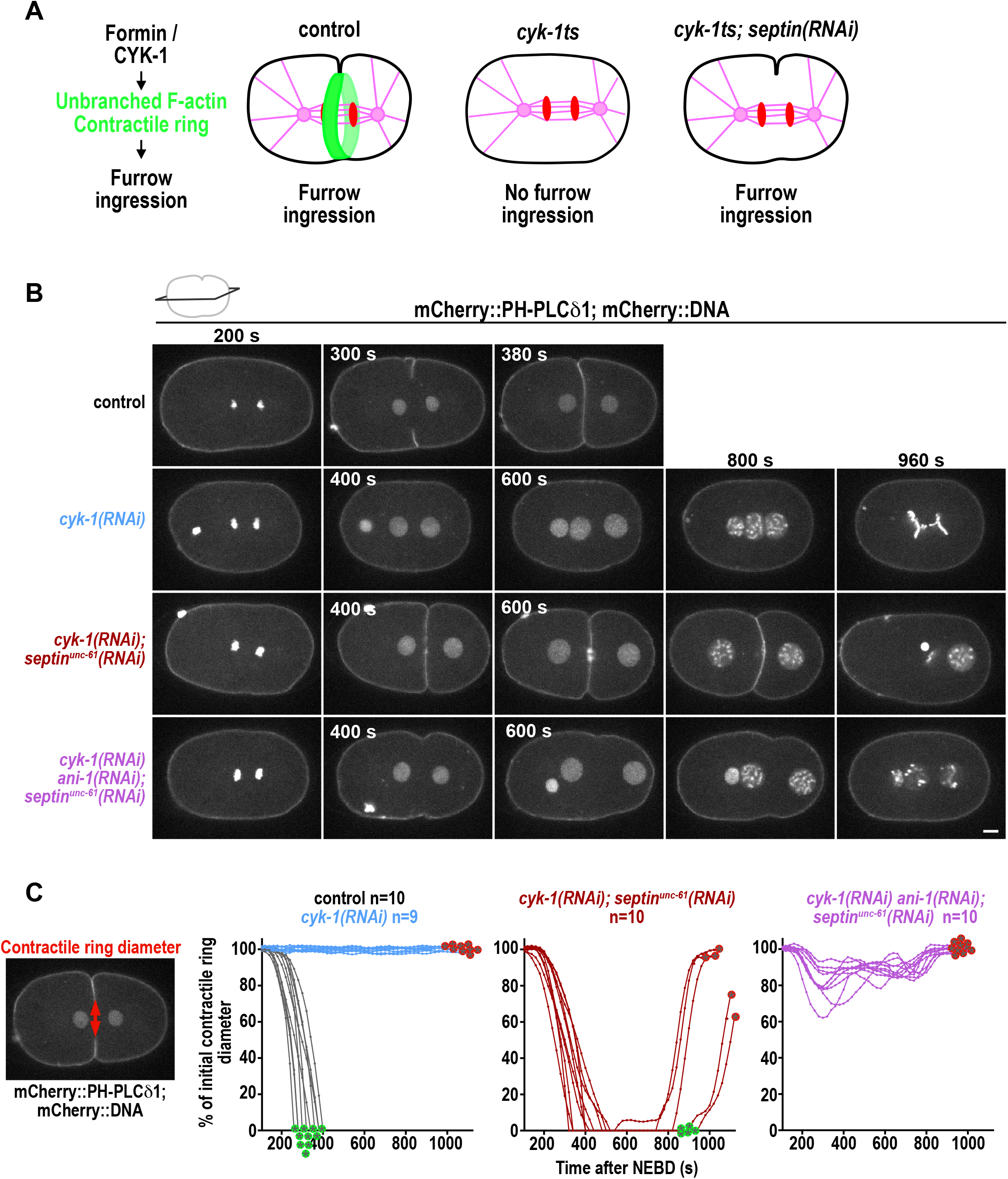
Furrow ingression of *cyk-1(RNAi); septin^unc-61^ (RNAi)* embryos requires ANI-1. **A)** During cytokinesis the formin CYK-1 polymerizes the unbranched F-actin of the contractile ring. In *cyk-1ts* mutants no contractile ring forms and cytokinesis fails, but, surprisingly furrow ingression was restored by depleting septin (Jordan *et al*., 2016). **B)** Central plane images of embryos expressing the membrane marker mCherry::PH-PLCδ1 and mCherry::DNA (Histone-H2B) and treated with the indicated RNAi conditions. Time is indicated in seconds (s) after NEBD. Scale bar is 5 μm. **C)** The contractile ring diameter is plotted as % of the initial contractile ring diameter over time for individual embryos for indicated RNAi conditions. Green and red encircled stars indicate whether embryos succeed or fail cytokinesis, respectively. n=number of embryos analyzed.

### ANI-1 localizes to septin-independent linear structures, especially after the depletion of the formin CYK-1

Since ANI-1 was required for furrow ingression, we analyzed ANI-1 localization by imaging endogenously NeonGreen-tagged ANI-1 on the cell cortex. In control anaphase embryos ANI-1 was enriched in cortical patches as previously published (Lewellyn *et al*., 2010), but also localized in linear structures, which sometimes radiate out of cortical patches (Fig. 2A, Movie 2). Comparison of the localization of ANI-1 with endogenously-tagged myosin II (NMY-2::mKate) revealed that NMY-2 was also present in the cortical patches (Werner et al., 2007) and occasionally decorated the linear ANI-1 structures. In CYK-1-depleted embryos cortical ANI-1 patches were absent and numerous linear structures, decorated with NMY-2, formed during early anaphase (Fig. 2A, Movie 2). To quantify the number of the linear ANI-1 structures, we measured the length and width of all structures in a defined region at the cell equator. Structures with a length/width ratio < 4 were assigned to the non-linear and ≥4 to the linear class (Fig. S3A). Linear ANI-1 structures were occasionally observed in control embryos and after *cyk-1(RNAi)* their number increased (Fig. 2B). In *cyk-1(RNAi)* linear ANI-1 structures had a length of ~2 - 9 μm and width of ~0.5 μm. To determine whether the overall ANI-1 distribution was altered after *cyk-1(RNAi)*, ANI-1 fluorescence intensity was measured along the cell cortex 180 s after NEBD, which represent the time point of contractile ring assembly. After *cyk-1(RNAi)*, ANI-1 was still enriched in an equatorial zone, although its intensity was slightly reduced in comparison to control embryos (Fig. S3B). After the co-depletion of septin^UNC-61^ and CYK-1, linear ANI-1 structures were still present, although cortical ANI-1 levels decreased further (Fig. 2A, B, S3B), consistent with the fact that septin and anillin interact in other organisms (Straight *et al*., 2005). Measurement of the orientation of ANI-1 at the cell equator during anaphase revealed that ANI-1 circumferentially aligns around the cell equator similar to F-actin in control embryos (Fig. 2C). Reduction of unbranched F-actin alone or together with septin^UNC-61^ did not prevent alignment of ANI-1 (Fig. 2C). Since ANI-1 appeared to form those linear structures, we determined whether septin also localized to linear structures after *cyk-1(RNAi)*. Indeed, we observed that after *cyk-1(RNAi)* endogenously-tagged septin^UNC-59^ was also present in linear structures similar to ANI-1 and septin^UNC-59^ cortical intensity levels were increased (Fig. S3C-E).

**Figure 2.**
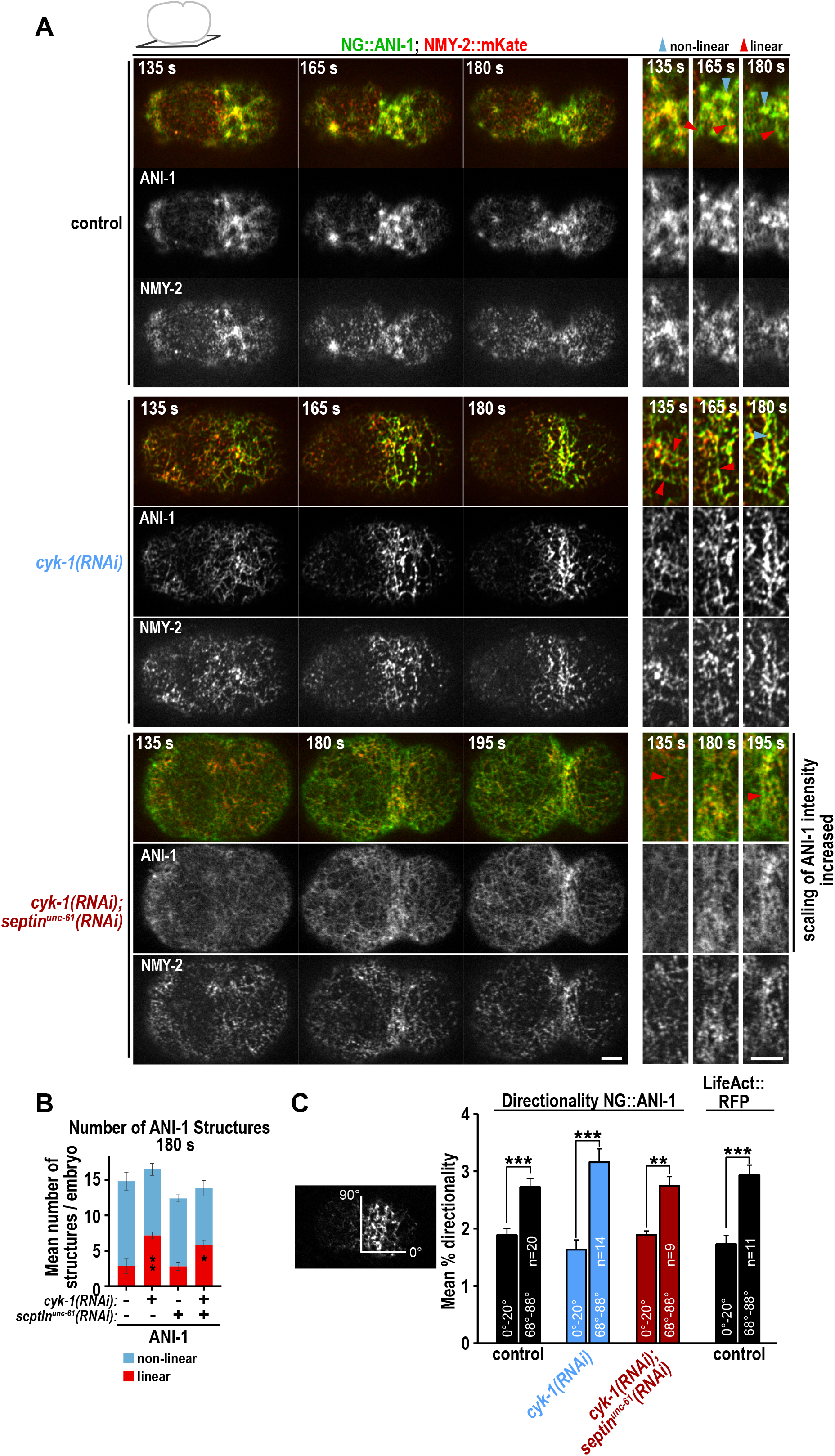
ANI-1 forms linear structures decorated by NMY-2 foci when CYK-1 formin nucleated F-actin levels are low. **A)** Confocal single z-plane images of the cell cortex of endogenously tagged NG::ANI-1 (green) and NMY-2::mKate (red) for indicated RNAi conditions during indicated time points after NEBD. Magnification of the equatorial region of the same embryos are shown to highlight the different structures NG::ANI-1 and NMY-2::mKate localize to. Since NG::ANI-1 levels are reduced after septin^UNC-61^ depletion (Fig. S3B), intensity scaling was increased to visualize the ANI-1 structures after septin co-depletion. Linear structures are highlighted by red arrowheads and non-linear by blue arrowheads. **B)** The mean number of linear and non-linear NG::ANI-1 structures per embryo at an equatorial region for the indicated RNAi conditions at 180 s after NEBD. P-values were calculated using Mann-Whitney or student *t*-test and represent *p<0.05 and **p<0.01 in comparison to control embryos treated without RNAi, n≥5 embryos for each condition. **C)** Mean percentage of 0-20° (anterior to posterior) and 68-88° (circumferentially) directionality measured for NG::ANI-1 and LifeAct::RFP at 180 s after NEBD for indicated RNAi conditions. Error bars are SEM and *P*-values were determined with student *t*-test and represent *** p<0.001; ** p<0.01. All scale bars are 5 μm. Error bars are SEM and n=number of embryos analyzed.

Since reduction of CYK-1-nucleated unbranched F-actin altered ANI-1 localization, we analyzed whether also Arp2/3-nucleated branched F-actin has an influence on ANI-1 by depleting ARX-2. Consistent with previous reports we observed a reduction of F-actin foci after *arx-2(RNAi)* (Fig. S4A) and a delay in furrow formation and ANI-1 accumulation (Fig. S4B, C) (Chan *et al*., 2018; Davies *et al*., 2014; Shivas and Skop, 2012). However, ARX-2 depletion did not impact ANI-1 localization to linear structures in control or *cyk-1(RNAi)* embryos (Fig. S4B, D, Movie 3). In summary, CYK-1-nucleated unbranched, but not Arp2/3-nucleated branched F-actin limits the formation of linear ANI-1 structures.

### NMY-2 is required for the orientation of the linear ANI-1 structures and furrow ingression in embryos depleted of CYK-1 and septin

We found that ANI-1 forms linear structures decorated by NMY-2 and supports furrow ingression in *cyk-1(RNAi); septin^unc-61^ (RNAi)* embryos. To test whether NMY-2 is required for the formation or alignment of the linear ANI-1 structure, we depleted NMY-2 together with CYK-1 or CYK-1 and septin^UNC-61^. Immunoblot analysis and quantification of cortical NMY-2 fluorescence intensity confirmed that *nmy-2(RNAi)* strongly reduced endogenously-tagged NMY-2 protein levels in worm lysates and on the cell cortex during anaphase in all tested RNAi conditions (Fig. S5A-C). Depletion of NMY-2 alone or in combination with CYK-1 or CYK-1 and septin^UNC-61^ did not prevent linear ANI-1 structure formation, although they were no longer circumferentially aligned around the cell equator (Fig. 3A-C, Movie 4). This suggests that NMY-2 aligns linear ANI-1 structures, perhaps by directly binding to the ANI-1 N-terminus. In normal cytokinesis myosin II binds and aligns F-actin and mediates furrow ingression (Osório et al., 2019), so we asked whether NMY-2 still mediates furrow ingression in *cyk-1(RNAi); septin^unc-61^(RNAi)* embryos. NMY-2 co-depletion abolished furrow ingression in *cyk-1(RNAi); septin^unc-61^(RNAi)* embryos (Fig. 3D, S5E), although ANI-1 was still enriched at the cell equator (Fig. 3A, S5D). In conclusion, NMY-2 is absolutely required to mediate furrow ingression after the reduction of CYK-1 and septins. Further, NMY-2 is involved in aligning the linear ANI-1 structures circumferentially but is not essential for their formation.

**Figure 3.**
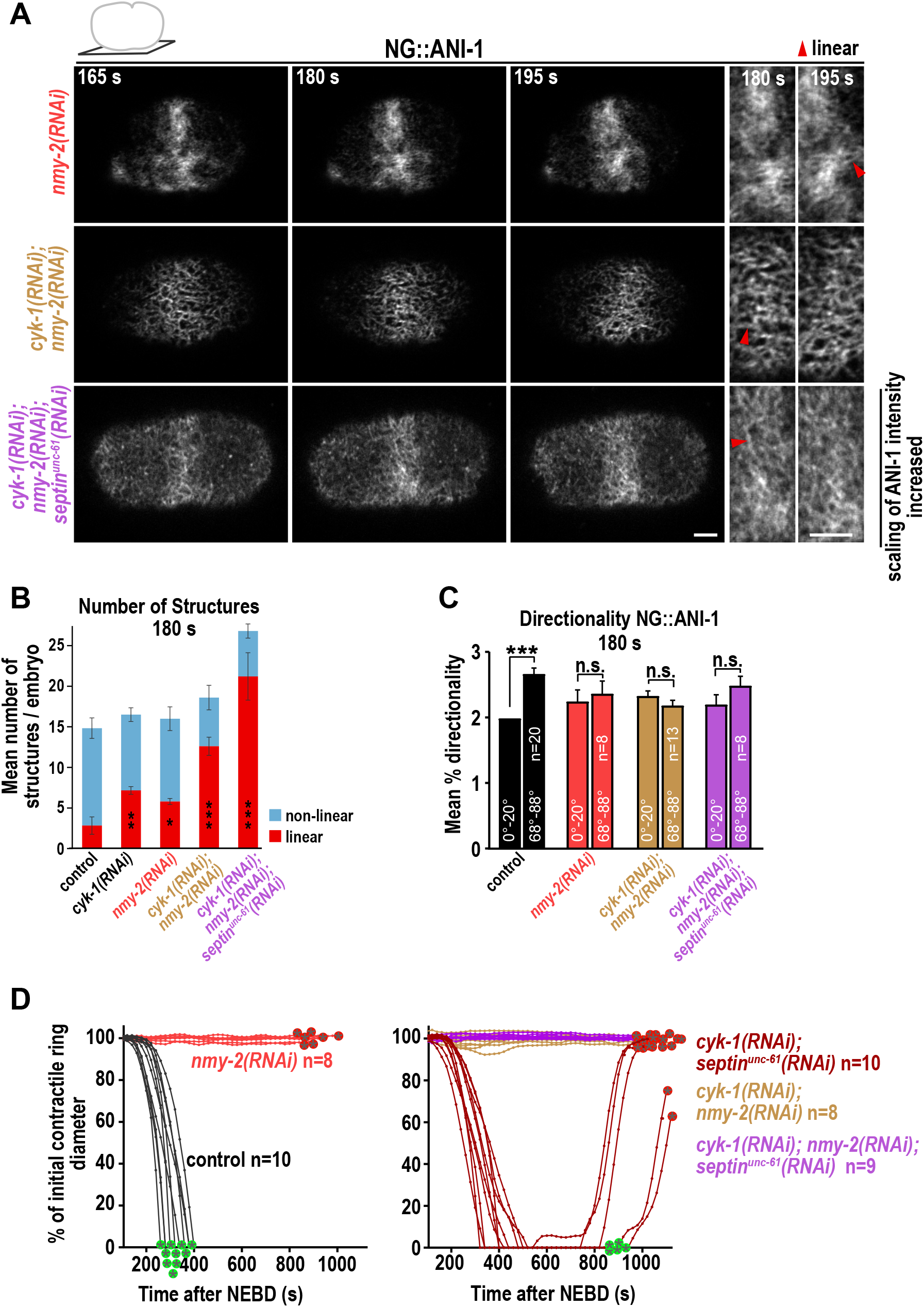
NMY-2 is not required for the formation of the linear NG::ANI-1 structures but for their circumferential alignment. **A)** Single z-plane cortical confocal images of NG::ANI-1 for the indicated RNAi conditions and time points after NEBD. Magnification of the cell equator is shown on the right. Since NG::ANI-1 levels are reduced after septin^UNC-61^ depletion, intensity scaling was increased to visualize the NG::ANI-1 structures after *septin^unc-61^(RNAi)*. Scale bars are 5 μm. **B)** Mean number of NG::ANI-1 structures at the cell equator per embryo for indicated RNAi conditions 180 s after NEBD, error bars are SEM, *P-* values were calculated using Mann-Whitney or student *t*-test and represent * p<0.05, ** p<0.01, *** p<0.001 in comparison to control embryos, n≥5 embryos for each condition. Control and *cyk-1(RNAi)* conditions are reproduced from Fig. 2B. **C)** Mean percentage of 0-20° (anterior to posterior) and 68-88° (circumferentially) directionality for NG::ANI-1 at indicated RNAi conditions 180 s after NEBD. Error bars are SEM and *P*-values were determined with student *t*-test and represent ***p<0.001; n.s. p>0.05. Control condition is reproduced from Fig. 2C. **D)** Plotted is the contractile ring diameter of each embryo over time for indicated RNAi conditions, n=number of embryos analyzed. Green and red encircled stars indicate whether embryos succeed or fail cytokinesis, respectively. Control and *cyk-1(RNAi); septin^unc-61^ (RNAi)* conditions are reproduced from Fig. 1C. For all n=number of embryos analyzed.

### ANI-1 and NMY-2 form membrane-localized structures in the absence of F-actin, independently of the ANI-1 N-terminus

After depolymerization of F-actin by Latrunculin A treatment in *Drosophila* cells, anillin and the RLC of myosin II enrich in circular foci that evolve into linear structures and these structures also form independently of each other (D’Avino *et al*., 2008; Hickson and O’Farrell, 2008; Oegema *et al*., 2000). In the *C. elegans* zygote, NMY-2 also forms circular foci after Latrunculin A addition (Michaux et al., 2018; Sobral et al., 2021). However, whether ANI-1 is also present in those circular NMY-2 foci and whether linear structures still form after F-actin depolymerization has not been investigated in *C. elegans*. To study this, we treated embryos with Latrunculin A, which caused the disappearance of F-actin from the cortex within ~80 sec (Fig. S6A-C) and the appearance of highly dynamic NMY-2 and ANI-1 structures that formed and disappeared quickly (Fig. S6D, E, Movie 5). After loss of cortical F-actin, NMY-2 and ANI-1 co-localized in numerous non-linear structures 180 s after NEBD that became linear at later anaphase time points (Fig. 4A, B). Measurement of fluorescence intensities at the cell cortex revealed that upon F-actin depolymerization NMY-2 and ANI-1 accumulated in an equatorial zone similar to control embryos, although NMY-2 fluorescence was increased at 255 s (Fig. S6F).

**Figure 4.**
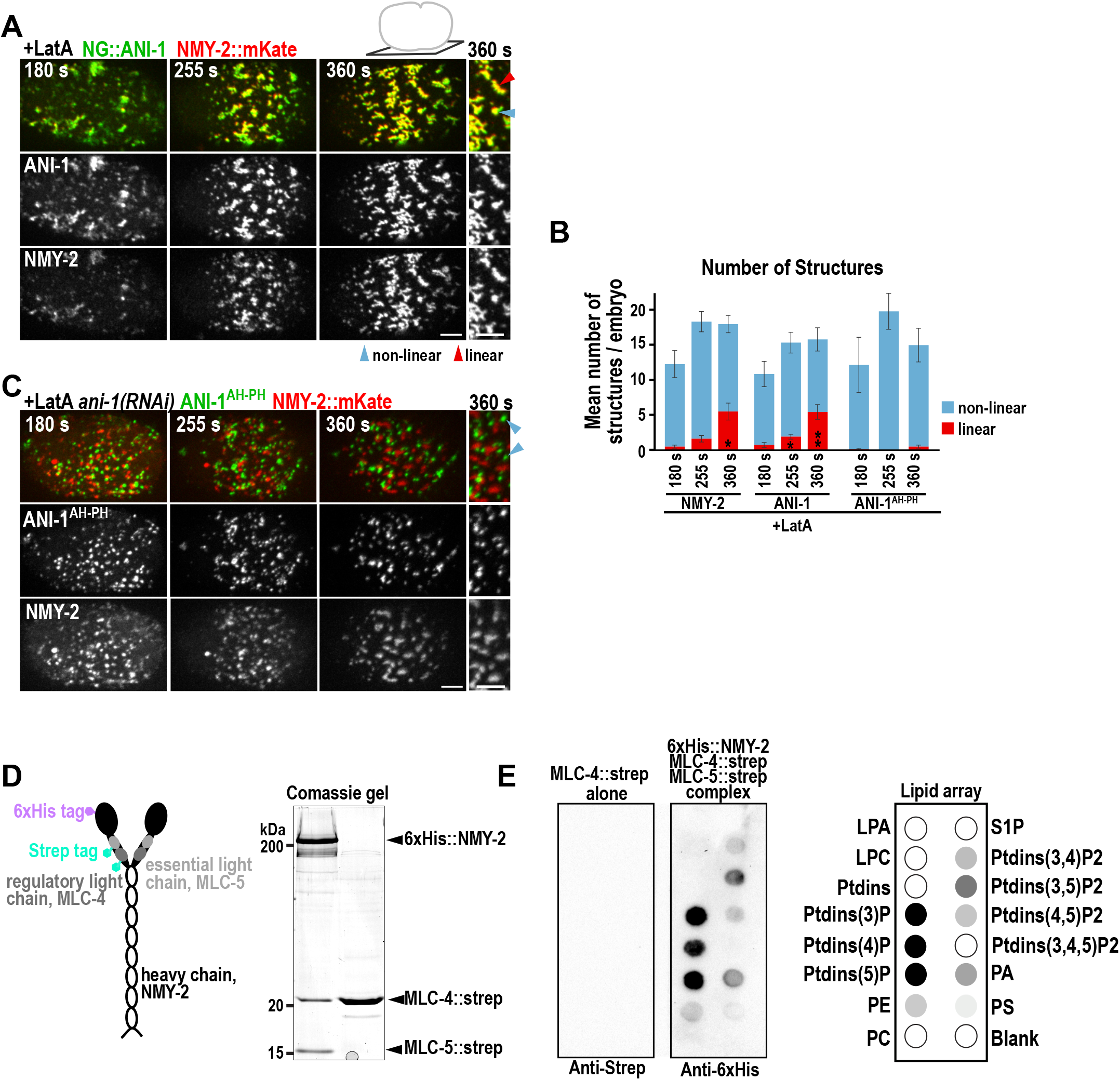
NMY-2 forms membrane localized structures independently of F-actin and the ANI-1 N-terminus and interacts with lipids *in vitro*. **A)** Maximum intensity projections of 10 cortical z-planes of permeabilized one-cell embryos treated with Latrunculin A (+LatA) expressing NG::ANI-1 (green) and NMY-2::mKate (red). A magnification of the cell equator is shown on the right and non-linear and linear structures are highlighted by blue and red arrowheads, respectively. **B)** Mean number of NMY-2::mKate, NG::ANI-1 and ANI-1^AH-PH^ structures at the cell equator per embryo for the indicated conditions. Error bars are SEM, *P*-values were calculated using Mann-Whitney or student *t*-test and represent * p<0.05, ** p<0.01 in comparison to 180 s, n≥5 embryos for each condition. **C)** Maximum intensity projections of 10 cortical z-planes of permeabilized embryos expressing ANI-1^AH-PH^ (green) (Tse et al., 2012) and NMY-2::mKate (red) treated with Latrunculin A and *ani-1(RNAi)*. A magnification of the cell equator is shown on the right and non-linear structures are highlighted by red arrowheads, respectively. **D)** Strep-tagged MLC-4 alone or in complex with 6xHis-tagged NMY-2, Strep-tagged MLC-5 were purified from insect cells and analyzed on a comassie-stained SDS-gel. **E)** Purified MLC-4 alone or in complex with 6xHis-tagged NMY-2 and Strep-tagged MLC-5 were incubated with lipid arrays and blotted with anti-Strep or anti-6xHis antibodies (left). Schematics of the lipid array and obtain results are shown on the right. Lipid array: LPA-lysophosphatidic acid; LPC - Lysophosphatidylcholines; PtdIns - Phosphatidylinositol; PE - Phosphatidylethanolamine; PC - Phosphatidylcholines; S1P - Sphingosine-1-phosphate. triglyceride (TG); PA – Phosphatidic acid; PS - Phosphatidylserine. All scale bars are 5 μm and time is display in sec after NEBD.

Vertebrate anillin binds myosin II with its N-terminus (Straight *et al*., 2005) and the colocalization of anillin and myosin II in foci after Latrunculin A treatment requires the anillin N-terminus in *Drosophila* cells (Carim et al., 2020). Since NMY-2 and ANI-1 structures co-localize after Latrunculin A treatment and NMY-2 is expected to interact with the ANI-1 N-terminus, we analyzed whether ANI-1 and NMY-2 still form structures that co-localize when the N-terminus of ANI-1 is absent. We used an ANI-1 fragment that comprises the RBD, C2 and PH domain (Δ1-680, ANI-1^AH-PH^ (Tse et al., 2012)) and depleted endogenous ANI-1 by RNAi to prevent NMY-2 binding via full-length endogenous ANI-1. After Latrunculin A addition NMY-2 and ANI-1^AH-PH^ both formed circular structures, but these did not co-localize and did not become linear over time (Fig. 4B, C, Movie 6). This shows that the N-terminus of ANI-1 is not required for the formation of non-linear membrane structures but is essential for the co-localization with NMY-2 structures. Thus, like in other organisms the N-terminus of ANI-1 possibly interacts with NMY-2.

Since NMY-2 formed membrane-localized structures independently of its binding partners F-actin and of the ANI-1 N-terminus, we hypothesized that NMY-2 could bind the membrane directly after its activation by RhoA signaling. To test this, we purified NMY-2 in a complex with the myosin essential light chain 4 (MLC-4) and the myosin regulatory light chain (MLC-5). We found that the NMY-2/MLC-4/MLC-5 complex, but not MLC-4 alone, bound to several lipids known to be enriched in the plasma membrane (Fig. 4D, E). This suggests that the NMY-2/MLC-4/MLC-5 complex directly binds the plasma membrane.

In summary, in the absence of F-actin, ANI-1 and NMY-2 co-localize to non-linear and linear structures, although the formation of linear structures takes much longer than in *cyk-1(RNAi)*. The co-localization but not the formation of the ANI-1 and NMY-2 non-linear structures requires the ANI-1 N-terminus, suggesting that the binding of NMY-2 to the ANI-1 N-terminus pulls those structures together. Since the NMY-2/MLC-4/MLC-5 complex bound to several membrane lipids, we conclude that this complex represents an independent membrane anchor.

### The linker region between the N- and C-terminus of ANI-1 is essential for linear structure formation

ANI-1 contains multiple distinct domains: an actin-binding domain (ABD) (Tian *et al*., 2015) and a predicted myosin II binding domain (MBD) (Maddox *et al*., 2005) at the N-terminus, and a RBD, a C2 membrane binding and a PH domain at the C-terminus (Sun *et al*., 2015) (Fig. S7A). Sequence analysis predicts that the ANI-1 N-terminus contains numerous intrinsically disordered regions (IDRs) similar to human anillin (Fig. S7A, B) and yeast Mid1p (Chatterjee and Pollard, 2019).

To analyze whether the N- or C-terminal halves of ANI-1 form linear structures, we generated animals expressing transgenic ANI-1^C-term^ (ANI-1 Δ48-680), which comprises the RBD, the C2 and the PH domain, or ANI-1^N-term^ (ANI-1 Δ765-1159), which comprises the putative MBD, the ABD and the linker region. Since the N-terminal half lacks a membrane binding motif we fused it with the membrane binding regions of RHO-1 (RhoA in *C. elegans*, Fig. S8A). To test the function and localization of the ANI-1 variants in the absence of endogenous ANI-1, we rendered the transgenes *ani-1* RNAi resistant so that we could specifically deplete endogenous ANI-1 by RNAi (Fig. S8B).

As a control we generated a full-length wild-type ANI-1 transgene (ANI-1^WT^), which rescued embryonic lethality after *ani-1(RNAi)* suggesting that it is functional (Fig. S8C). ANI-1^WT^ was enriched at the cell equator and after depletion of CYK-1 and co-depletion of CYK-1 and septin^UNC-61^, the number of linear ANI-1^WT^ structures was increased and they circumferentially aligned around the cell equator (Fig. 5A-F, Movie 7-9). Together, the GFP-tagged ANI-1^WT^ transgene behaves similar to endogenously tagged NG::ANI-1.

**Figure 5.**
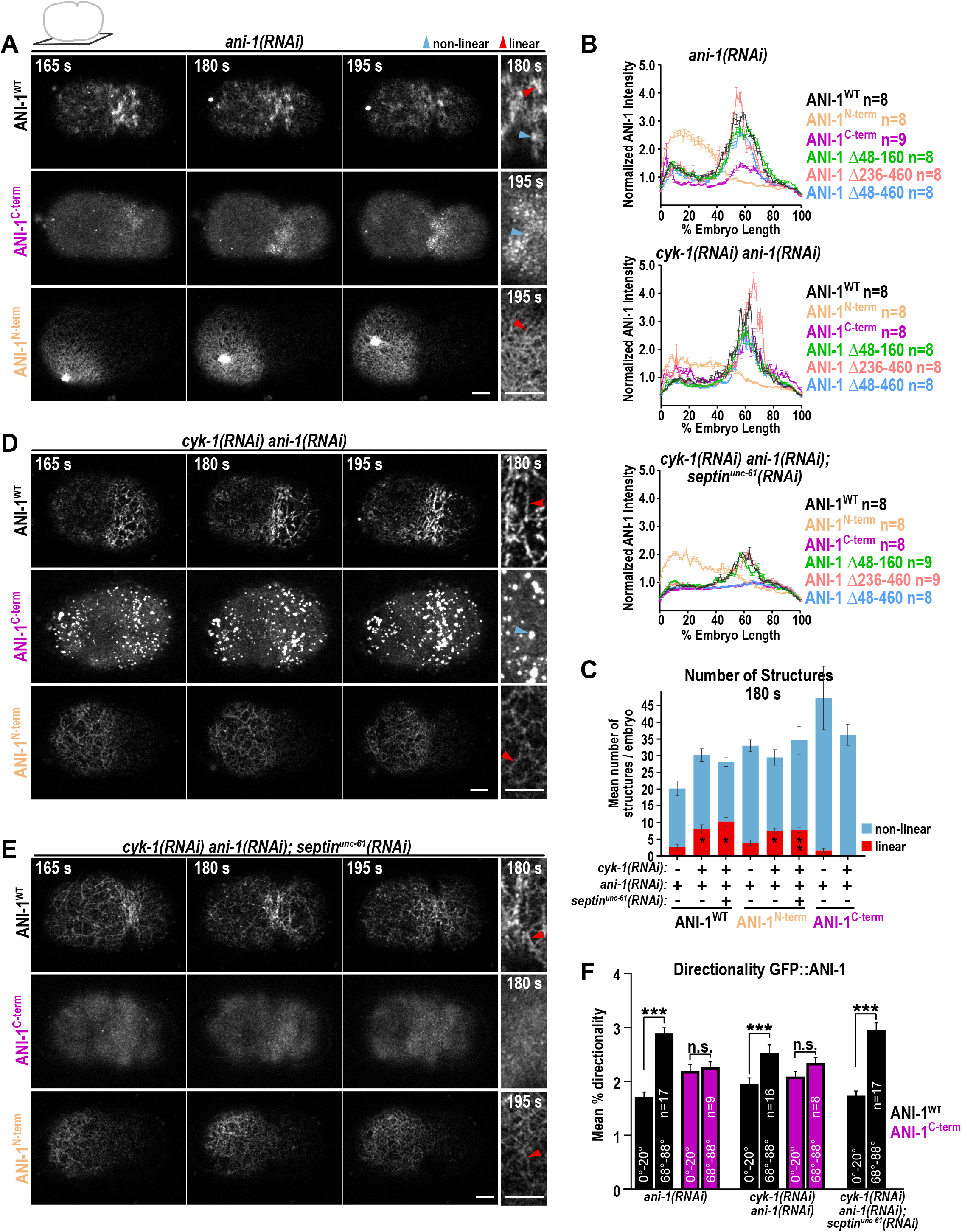
The ANI-1^N-term^ forms linear and the ANI-1^C-term^ circular structures. **A, D, E)** Confocal cortical single z-plane images of GFP-tagged ANI-1^WT^, ANI-1^N-term^ and ANI-1^C-term^ expressing embryos for the indicated RNAi conditions and time points after NEBD. Magnification of an equatorial (ANI-1^WT^, ANI-1^C-term^) and anterior (ANI-1^N-term^) region are shown on the right. Linear structures are highlighted by red and non-linear by blue arrowheads. To better visualize the localization of ANI-1^N-term^ after *cyk-1(RNAi) ani-1(RNAi)*(D) and for all transgenes after septin co-depletion (E), the signal intensity scaling was increased in those conditions. Scale bar is 5 μm. **B)** Mean normalized cortical fluorescence intensity of indicated GFP-tagged ANI-1 variants from the anterior to the posterior cortex for indicated RNAi conditions 180 s after NEBD. **C)** Mean number of structures per embryo at the cell equator for the different GFP-tagged ANI-1 proteins after RNAi depletions 180 s after NEBD. Note: for ANI-1^N-term^ the structures were counted in a region located at the anterior cortex. P-values were calculated using Mann-Whitney or student *t*-test and represent * p<0.05, ** p<0.01 in comparison to *ani-1(RNAi)* embryos expressing the same transgene, n≥5 embryos for each condition. **F)** Mean percentage of 0-20° (anterior to posterior) and 68-88° (circumferentially) directionality measured for GFP-tagged ANI-1 proteins for indicated RNAi conditions 180 s after NEBD. *P*-values were determined with student *t*-test and represent *** p<0.001; n.s. p>0.05. Error bars are SEM and n=number of embryos analyzed.

ANI-1^N-term^ or ANI-1^C-term^ were found to express at similar levels as ANI-1^WT^ and both mutants partially rescued embryonic lethality in the absence of endogenous ANI-1 (Fig. S8C, D). The ANI-1^C-term^ fragment was enriched at the cell equator, but its cortical levels were reduced in comparison to those of ANI-1^WT^ (Fig. 5A, B). ANI-1^C-term^ formed very few linear but numerous non-linear structures (Fig. 5A, C, Movie 7). After CYK-1 depletion, ANI-1^C-term^ did not form any linear structures but only non-linear ones that did not show circumferential alignment (Fig. 5C, D, F, Movie 8). Co-depletion of CYK-1 and septin^UNC-61^ resulted in a strong reduction of cortical ANI-1^C-term^ levels in comparison to ANI-1^WT^ (Fig. 5B, E, Movie 9) and the absence of non-linear foci, suggesting that its enrichment in non-linear structures requires septin binding. However, transient pulsatile accumulation of ANI-1^C-term^ was still observed, possibly due to active RhoA binding (Movie 9). Since almost no ANI-1^C-term^ localized to the cell equator after *cyk-1(RNAi); septin^unc-61^(RNAi)*, we did not perform the structure or directionality analysis. The ANI-1^N-term^ fragment was enriched at the anterior cortex (Fig. 5A, B, Movie 12), probably because RhoA and septin binding domains were missing. Therefore, we performed the structure analysis of the ANI-1^N-term^ fragment in a region of the anterior cortex and found that it formed linear structures in all RNAi conditions (Fig. 5A, C, D, E, Movie 7-9). Together this demonstrates that the N-terminal but not the C-terminal half of ANI-1 is sufficient to form linear structures after *cyk-1(RNAi)*.

Since the N-terminus was sufficient for linear structure formation, we tested whether the predicted MBD and ABD domains are required for their formation. We generated ANI-1 transgenes where the MBD (Δ48-160, Fig. S8A) or the ABD (Δ236-460) or both (Δ48-460) domains had been deleted. All transgenes were expressed at similar levels to ANI-1^WT^ and rescued embryonic lethality after *ani-1(RNAi)* (Fig. S8C, D). The cortical fluorescence intensities, the number of linear structures and the circumferential alignment of the Δ48-160, Δ236-460 and Δ48-460 ANI-1 mutants were comparable to that of ANI-1^WT^ after *ani-1(RNAi)* or *cyk-1(RNAi) ani-1(RNAi)* (Fig. 5B, S9A-E, Movie 10 and 11). After co-depletion of CYK-1 and septin^UNC-61^, the cortical levels of ANI-1 Δ48-460 were strongly reduced at the cell equator, but the levels of ANI-1 Δ48-160 and ANI-1 Δ236-460 were comparable to ANI-1^WT^. In those embryos ANI-1 Δ48-160 and ANI-1 Δ236-460 still formed linear structures that aligned circumferentially around the cell equator (Fig. 5B, S9C-E, Movie 12). Together, our ANI-1 mutant analysis reveals that neither the putative MDB nor the ABD, but rather the linker region is essential for linear structure formation and their circumferential alignment after CYK-1 depletion.

### The ANI-1^N-term^ partially recues furrow ingression in *cyk-1(RNAi); septin^unc-61^ (RNAi)* embryos

To investigate whether the different ANI-1 mutants support furrow ingression, we filmed embryos in the central plane after co-depletion of CYK-1 or CYK-1 and septin^UNC-61^. As expected, depletion of CYK-1 in ANI-1^WT^ expressing embryos prevented furrow ingression and co-depletion of CYK-1 and septin^UNC-61^ resulted in complete furrow ingression (Fig. 6). ANI-1 Δ48-160 and ANI-1 Δ236-460 rescued cleavage furrow ingression similar to ANI-1^WT^ (Fig. 6) consistent with the fact that they formed linear structures. ANI-1 Δ48-460 and ANI-1^C-term^ did not rescue furrow ingression, but as both localized only weakly to the cortex no rescue was expected (Fig. S10). Importantly, five out of eight embryos expressing ANI-1^N-term^ fully ingressed a cleavage furrow and in two of those embryos the cleavage furrow remained closed until the end of the movie. This demonstrated that the N-terminus of ANI-1, even though not able to fully enrich at the cell equator, is able to partially rescue the cytokinesis failure in *cyk-1(RNAi) ani-1(RNAi); septin^unc-61^ (RNAi)* embryos.

**Figure 6.**
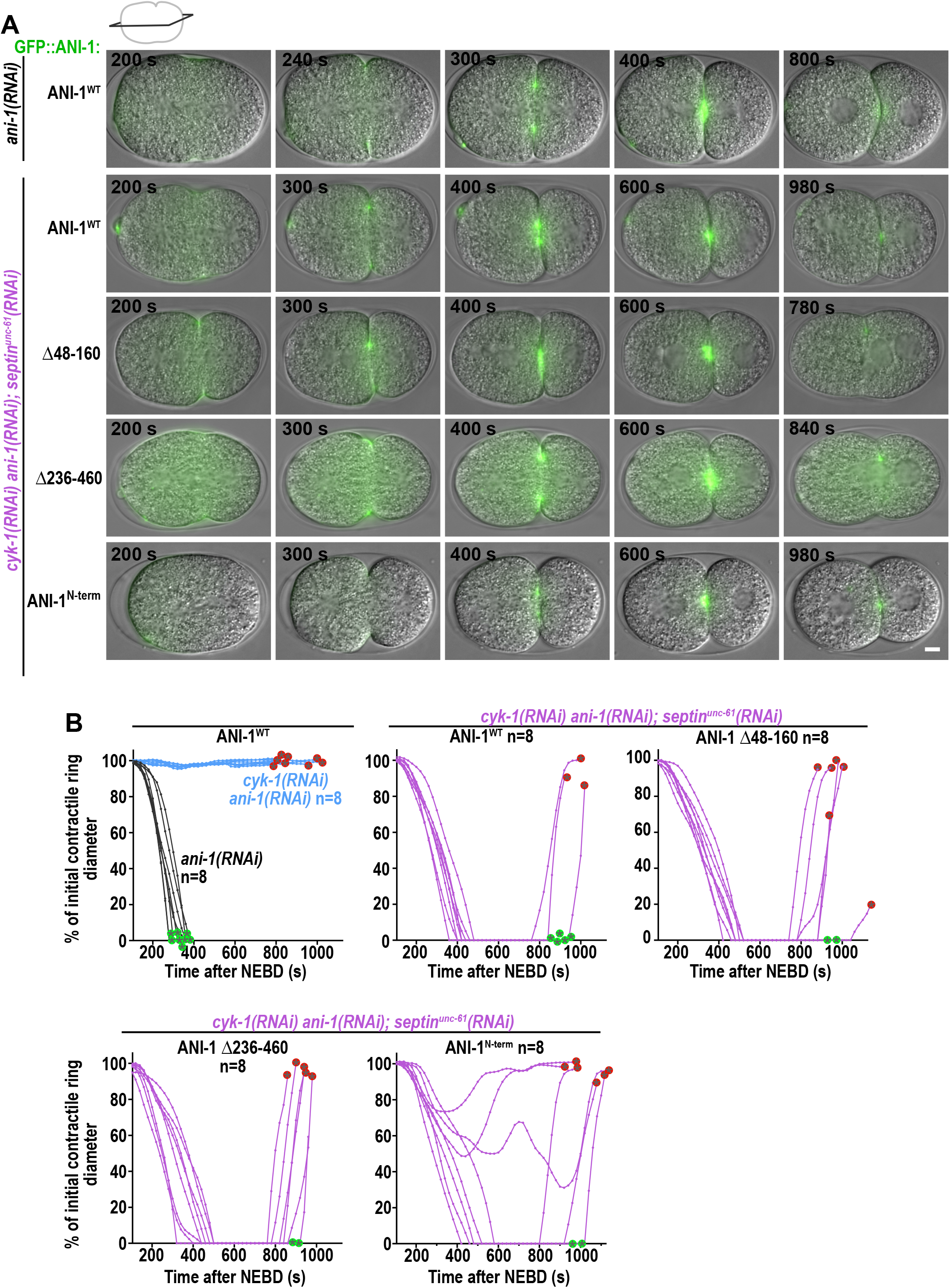
The N-terminal ANI-1 fragment partially rescues furrow ingression in *cyk-1(RNAi) ani-1(RNAi); septin^unc-61^(RNAi)* **A)** Merged DIC and wide-field fluorescent images at the central plane for the different GFP-tagged ANI-1 mutant proteins for the indicated RNAi conditions and time points after NEBD. Scale bar is 5 μm. **B)** Plotted is the contractile ring diameter of each embryo over time for indicated RNAi conditions and GFP-tagged ANI-1 mutants, n=number of embryos analyzed. Green and red encircled stars indicate whether embryos succeed or fail cytokinesis, respectively.

## Discussion

In animal cells the contractile ring consists of formin-nucleated unbranched F-actin, which circumferentially aligns around the cell equator. Myosin-dependent constriction of the contractile ring mediates cleavage furrow ingression (Leite et al., 2019). Here we discover an ANI-1-dependent mechanism that promotes furrow ingression when the levels of formin-nucleated unbranched F-actin and septins are reduced (Fig. S11). In this situation ANI-1 forms a meshwork of linear structures decorated by NMY-2 that circumferentially align around the cell equator. Based on previous reports and our data we propose the following model for furrow ingression after the depletion of septins and formins: active RHO-1 recruits ANI-1 and NMY-2 to the cell equator in anaphase. ANI-1 is tethered to the equatorial membrane by its C2 and RBD domain. The disordered ANI-1 N-terminus extends into the cytoplasm, interacts with neighboring ANI-1 molecules and forms the linear structures. Membrane-bound NMY-2 minifilaments crosslink and circumferentially align linear ANI-1 structures around the cell equator. Adjacent linear ANI-1 structures interact and shorten in length. This shortening of linear ANI-1 structures is propagated around the ANI-1 network by the crosslinker NMY-2. Since both ANI-1 and NMY-2 are linked to the membrane the contraction force is conveyed to the membrane and the furrow forms and ingresses. Some linear ANI-1 structures are present in wild type embryos and could also assist with furrow ingression by a similar mechanism in the presence of normal levels of formin-nucleated F-actin.

### NMY-2 is an independent membrane anchor, which links the actin cortex with the plasma membrane

NMY-2 enriches in foci on the equatorial plasma membrane after F-actin depolymerization even in the absence of the ANI-1 N-terminus. Furthermore, our *in vitro* binding assays show, that NMY-2, together with the RLC and ELC, interacts with multiple plasma membrane lipids. Together, our *in vivo* and *in vitro* data suggest that NMY-2, after activation by RhoA signaling, binds the membrane directly. Membrane binding of myosin II, seems conserved in other organisms as in *Drosophila* cells myosin also localizes independently of anillin and F-actin to the membrane (Carim *et al*., 2020). In fission yeast Mid1p targets Myo2 to the cytokinesis nodes (Motegi et al., 2004; Saha and Pollard, 2012), and Mid1p but not Myo2 leaves the nodes during ring constriction suggesting that Myo2 could also bind the plasma membrane. A previous study found that myosin II interacts with lipids through its RLC binding site and lipid binding resulted in displacement of the RLC (Liu et al., 2016). This implied that membrane-bound myosin II is in the inactive state, since the RLC is required for myosin II activity. In contrast, we observed that NMY-2 interacts with membrane lipids in the presence of the RLC and ELC suggesting that NMY-2 can bind the membrane in its active state. Together, in *C. elegans*, membrane-bound active NMY-2 might represent an independent membrane anchor, which links the plasma membrane with the actin cortex and thereby could transmit forces generated in the contractile ring to the plasma membrane. NMY-2 membrane tethering might function redundantly with other membrane binding components such as ECT2 (Su et al., 2011), anillin (Liu *et al*., 2012), and RacGAP1 (Lekomtsev et al., 2012). Determining the region of the NMY-2/MLC-4/MLC-5 complex that interacts with the plasma membrane and finding out how membrane binding is coupled with NMY-2 activation are interesting questions to pursue in the future.

### Formation of linear ANI-1 structures is modulated by the amount of F-actin

We discover that ANI-1 not only localizes to cortical patches (Lewellyn *et al*., 2010; Maddox *et al*., 2005), but also to some linear structures during cytokinesis in the one-cell *C. elegans* embryo. After the reduction of formin-nucleated F-actin, numerous linear ANI-1 structures form and circumferentially align around the cell equator. How could F-actin limit the formation of linear ANI-1 structures? We speculate that when unbranched F-actin levels are high, ANI-1 binds F-actin and distributes along the filaments, which limits contacts between ANI-1 molecules or restrains extension or flexibility of the disordered ANI-1 N-terminus. When unbranched F-actin levels are low, such as in the case of CYK-1 depletion, ANI-1 N-termini are free, accessible to self-interact and form linear structures. In the complete absence of F-actin (Latrunculin A treated embryos) ANI-1 first forms circular structures that become linear at late anaphase time points. Since linear ANI-1 structures were forming much slower in the presence of Latrunculin A, short actin filaments might bridge adjacent linear ANI-1 structures to accelerate the process. We did not observe a change in ANI-1 structure formation after the reduction of Arp2/3-generated branched F-actin suggesting that ANI-1 preferentially binds CYK-1-nucleated unbranched F-actin. Filamentous-type anillin was also observed along F-actin bundles in unperturbed vertebrate cells (Oegema *et al*., 2000), after myosin II inhibition (Straight *et al*., 2005) and in Latrunculin A treated *Drosophila* cells (Hickson and O’Farrell, 2008). Thus, the competence of anillin to form linear structures may be a conserved feature from worms to humans.

### The C-terminal half of ANI-1 concentrates at the cell equator in septin-dependent circular structures

Anillin’s C-terminal half harbors three membrane interaction elements: the RBD, the C2 and the PH domain, and each of them contributes to membrane localization of anillin in human cells (Liu *et al*., 2012; Piekny and Glotzer, 2008; Sun *et al*., 2015). Importantly, the ANI-1^C-term^ forms circular and not linear structures in *cyk-1(RNAi)* and Latrunculin A treated embryos. The localization of the ANI-1^C-term^ to the membrane strongly depends on septins, thus *C. elegans* ANI-1, similar to anillin in other organisms, should bind to septins via the PH domain (Kinoshita et al., 2002; Liu *et al*., 2012; Oegema *et al*., 2000; Piekny and Glotzer, 2008). Consistent with the fact that septins are not essential for the formation of linear ANI-1 structures, we observe that the ANI-1 N-terminal half, which lacks the predicted septin-binding region, is sufficient for linear structure formation. Since ANI-1 is reduced after septin depletion and septin recruitment to the contractile ring also depends on ANI-1 (Liu *et al*., 2012; Maddox *et al*., 2005), ANI-1 and septin stabilize each other on the plasma membrane.

After septin and CYK-1 co-depletion, a residual pulsatile membrane localization of the ANI-1^C-term^ persisted. This could reflect direct binding to the plasma membrane and/or binding to active RhoA. Given that the ANI-1^C-term^ membrane localization strongly depends on septins, changes in ANI-1^C-term^ dynamics cannot be exclusively attributed to active RHO-1 dynamics. Therefore, employing the ANI-1^C-term^ as a biosensor for active RHO-1 (Michaux *et al*., 2018; Reymann *et al*., 2016; Tse *et al*., 2011) should be done with caution.

### The disordered N-terminus is sufficient for linear structure formation

The membrane-targeted N-terminal ANI-1 half localized in linear structures. Further, our mutant analysis of full-length ANI-1 revealed that neither the MBD nor the ABD nor both domains together are required for ANI-1 linear structure formation in control and *cyk-1(RNAi)* embryos. Only further shortening and removal the linker region prevented the formation of linear structures and circular septin-dependent foci formed instead. Since the N-terminus of Mid1p forms high molecular weight oligomers (Celton-Morizur et al., 2004) one possibility is that the ANI-1 N-terminus assembles into linear structures by self-interaction mediated oligomerization. Furthermore, a liquid-liquid phase separation mechanisms might drive the formation of linear ANI-1 structures. Our structure prediction reveals that the N-terminus of ANI-1 is highly disordered but the C-terminus is not. This feature of ANI-1 is conserved from fission yeast (Chatterjee and Pollard, 2019) to worms and humans. Disordered regions frequently mediate the formation of biomolecular condensates by liquid-liquid phase separation (Ong and Torres, 2020) and indeed the disordered N-terminus of Mid1p forms condensates *in vitro* (Chatterjee and Pollard, 2019). Liquid droplets are typically isotropic but their shape and material properties are tunable in multiple ways. For example, isotropic liquid condensates of FUS transform into spindle-shaped condensates when F-actin is incorporated in the droplets (Scheff et al., 2020) or altering the concentration of the actin crosslinker filamin influences the shape and properties of F-actin droplets (Weirich et al., 2017). We speculate that linear ANI-1 structures could form by similar principles: the isotropic spherical shape of the biomolecular ANI-1 condensates is biased to an extended anisotropic form by additional factors such as remaining F-actin polymers, other proteins or posttranslational modifications of ANI-1. Single deletion of the N-terminal MBD and ABD or both together (ANI-1^Δ48-460^) did not cause embryonic lethality or alter ANI-1 localization in control and *cyk-1(RNAi)* embryos. Depletion of septins strongly reduced the cortical fluorescence intensity of the ANI-1^Δ48-460^ mutant. Thus, in addition to septins, the MBD and ABD redundantly contribute to equatorial enrichment. Similarly, in human and *Drosophila* cells the MBD and ABD of anillin are not essential but contribute to successful cytokinesis (Jananji *et al*., 2017; Piekny and Glotzer, 2008).

### The ANI-1 N-terminus promotes furrow ingression after co-depletion of formin and septins

During cytokinesis F-actin polymerization at the cell equator is induced by diaphanous-type formins and CYK-1 is the only diaphanous formin in *C. elegans* (Severson *et al*., 2002; Swan *et al*., 1998). Moreover, none of the other formins was found to act redundantly with CYK-1 during the division of early *C. elegans* blastomeres (Davies *et al*., 2018). In CYK-1-depleted embryos cleavage furrow ingression failed, but double depletion of CYK-1 and septins could restore furrow ingression similar to previous observations using a *cyk-1* temperature sensitive allele (Jordan *et al*., 2016). We found that after CYK-1 depletion, ANI-1 forms linear structures that circumferentially align around the cell equator and cortical septin^UNC-59^ levels increase. This indicates that septins or the excess thereof, hinder cleavage furrow ingression in CYK-1-depleted embryos. As septins bind ANI-1 and contribute to ANI-1 targeting to the cell equator, a reduction in septin levels might weaken the interaction between ANI-1 and the plasma membrane, and consequently lead to increased mobility of ANI-1, which could facilitate the interaction between ANI-1 molecules and the interaction between ANI-1 and NMY-2. We observed that all *cyk-1(RNAi); septin^unc-61^ (RNAi)* embryos fully ingress a cleavage furrow but half of them fail in abscission and the furrow regresses. Septins are essential for abscission in *Drosophila* and human cells (Estey *et al*., 2010; Kechad *et al*., 2012) but not in *C. elegans* embryos (Green *et al*., 2013). Thus, our data suggest a functional redundancy between unbranched F-actin and septins during abscission in *C. elegans*.

Our data suggest that the N-terminus of ANI-1 interacts with NMY-2, since co-localization of NMY-2 and ANI-1 after Latrunculin A treatment requires its presence. Consistent with that, NMY-2 was required for the circumferential alignment of the linear ANI-1 structures. Surprisingly, deletion of the putative MBD did not abrogate linear ANI-1 structure alignment. Thus, either the NMY-2 binding region is not properly predicted and is actually located in ANI-1’s ABD and/or linker region or NMY-2 does not align ANI-1 via direct binding. Since CYK-1-depleted embryos only exhibit short-range cortical flows (Naganathan et al., 2018), the alignment of the linear ANI-1 structures around the cell equator is unlikely to be flow-dependent, as suggested for unbranched F-actin (Reymann *et al*., 2016). Therefore we favor alternative models where localized motor-activity of NMY-2 and crosslinkers promotes equatorial circumferential alignment (Leite *et al*., 2020).

Furrow ingression was only supported by the ANI-1 fragments that still formed linear structures. Furthermore, those linear ANI-1 structures circumferentially align around the cell equator, similar to unbranched F-actin in control embryos. Therefore, it is tempting to speculate that those structures mediate furrowing in septins and CYK-1-depleted embryos. Linear ANI-1 structures could bridge the residual unbranched actin filaments so that NMY-2 can bind to them. Then, NMY-2 generates force as a motor or tension as a crosslinker by interacting with the remaining F-actin similar to what happens in normal contractile rings (Leite *et al*., 2019). However, since F-actin did not accumulate at the equator after CYK-1 and septin co-depletion and the ABD of ANI-1 was not required for furrow ingression, we favor an alternative model. We hypothesize that linear ANI-1 structures represent a novel type of network that contracts by biomolecular condensation. ANI-1 is predicted to have numerous IDRs at the N-terminus, and we showed that the N-terminus is sufficient for linear structure formation and furrow ingression. IDRs are frequently found in proteins that form biomolecular condensates by liquid-liquid phase separation. Recent reports demonstrate that biomolecular condensation can also exert forces on cellular structures and alter the shape of membranes (Ganar, 2021). Interactions between biomolecular condensates distributed along a DNA molecule pull the DNA molecule into the condensates and thereby apply force to the DNA polymer (Quail et al., 2021). The formation of biomolecular condensates of FUS on the surface of membrane vesicles generates compressive stress and causes inward membrane bending (Yuan et al., 2021). Strikingly, inward membrane bending was also observed when biomolecular condensates formed on the cytoplasmic surface of yeast cells and this mechanism contributes to actin-independent endocytosis (Bergeron-Sandoval et al., 2021). We speculate that condensation of linear ANI-1 structures results in the contraction of the ANI-1 network and ultimately in the inward bending of the plasma membrane (Fig. S11). The contractile force generated by ANI-1 condensation is then propagated around the ANI-1 network by NMY-2 filaments, which bind and crosslink the ANI-1 linear structures. As a first step, future work will need to show that *C. elegans* ANI-1, like Mid1p from fission yeast, also undergoes liquid-liquid phase separation, although we consider this to be highly likely due to the fact that the disordered disposition of the N-terminus is highly conserved (Chatterjee and Pollard, 2019). Linear ANI-1 structures are also observed in wild-type embryos that have normal F-actin levels. So, it is possible that ANI-1 linear structures may assist with furrow ingression in a normal situation and take over the process when unbranched F-actin levels are reduced.

In summary, we find that ANI-1 forms linear structures and supports furrow ingression when unbranched F-actin levels are low. Although actomyosin-dependent constriction is the most actively investigated mechanism for cytokinetic furrowing, alternative ways have been reported to exist. For example, furrow ingression is mediated in *Dictyostelium* and human RPE1 by the opposite migration direction of the two daughter cells (Dix *et al*., 2018; Nagasaki *et al*., 2009; Neujahr *et al*., 1997), and in archaea by supercoiling of the elastic ESCRT-III filament (Harker-Kirschneck et al., 2022; Tarrason Risa et al., 2020). Our findings highlight the existence of an additional mechanism of cleavage furrow ingression that is likely to contribute to the enormous robustness of cleavage furrow ingression during cytokinesis.

---

**Figure S1 Single, double and triple RNAi co-depletions of CYK-1 and septin are highly efficient**

**A)** Confocal single z-plane cortical images of endogenously tagged septin^UNC-59^::GFP for indicated RNAi conditions 180 s after NEBD.

**B)** Normalized septin^UNC-59^::GFP fluorescence intensity from the anterior (0% embryonic length) to the posterior (100%) cortex 180 s after NEBD. Graphs display the normalized mean septin^UNC-59^::GFP cortical fluorescence intensity for the indicated RNAi conditions.

**C)** Confocal cortical single z-plane images of endogenously tagged CYK-1::GFP (green) and LifeAct::RFP (red) expressing embryos for the indicated RNAi conditions 180 s after NEBD.

**D)** Normalized mean CYK-1::GFP (left) and LifeAct::RFP (right) fluorescence intensity from the anterior (0% embryonic length) to the posterior (100%) cortex 180 s after NEBD.

**E)** Immunoblot of septin^UNC-59^::GFP and CYK-1::GFP expressing worms with and without *septin^unc-59^(RNAi)* or *cyk-1(RNAi)* probed with antibodies against GFP and actin, as a loading control. The mean septin^UNC-59^::GFP or CYK-1::GFP protein levels (3 worm extracts) are indicated, star indicates a non-specific band.

All scale bars are 5 μm, error bars are SEM and n=number of embryos analyzed.

**Figure S2 ARX-2 depletion does not prevent furrow ingression in *cyk-1(RNAi); septin^unc-61^(RNAi)* embryos**

**A)** Central plane images of embryos expressing the membrane marker mCherry::PH-PLCδ1 and mCherry::DNA (Histone-H2B) and treated with the indicated RNAi conditions.

**B)** The contractile ring diameter is plotted as % of the initial contractile ring diameter over time for individual embryos for indicated RNAi conditions. Green and red encircled stars indicate whether embryos succeed or fail cytokinesis, respectively.

**C)** Immunoblot of NG::ANI-1 expressing worms with and without *ani-1(RNAi)* probed with antibodies against Flag and actin, as a loading control. The mean NG::ANI-1 protein levels (3 worm extracts) are indicated.

**D)** Confocal single z-plane images of the cell cortex of endogenously tagged NG::ANI-1 for indicated RNAi conditions at 180 s after NEBD.

**E)** Normalized NG::ANI-1 fluorescence intensity from the anterior (0% embryonic length) to the posterior (100%) cortex 180 s after NEBD for indicated RNAi conditions.

All scale bars are 5 μm, error bars are SEM and n=number of embryos analyzed.

**Figure S3 Equatorial septin^UNC-59^::GFP levels are elevated after CYK-1 depletion**

**A)** After filtering the original image, the length and width of each structure was measured in a region at the cell equator unless stated otherwise. Structures with a length/with ratio ≥4 were classified as linear and a length/with ratio <4 as non-linear.

**B)** Mean normalized NG::ANI-1 fluorescence intensity from the anterior to the posterior cortex at 180 s after NEBD for indicated RNAi conditions.

**C)** Confocal single z-plane images of embryos with endogenously GFP-tagged septin^UNC-59^ at selected time-points and RNAi conditions. Magnifications of the equatorial region at 165 s and 180 s after NEBD are shown.

**D)** Mean normalized septin^UNC-59^::GFP fluorescence intensity from the anterior to the posterior cortex for control and *cyk-1(RNAi)* treated embryos at 180 s after NEBD. Linear structures are highlighted by red arrowheads and non-linear by blue arrowheads.

**E)** Mean number of septin^UNC-59^::GFP structures per embryo for the indicated conditions at 180 s after NEBD and n≥5 embryos for each condition.

Error bars are SEM, all scale bars are 5 μm and n=number of embryos analyzed.

**Figure S4 ARX-2 depletion does not influence NG::ANI-1 localization**

**A)** Confocal single z-plane images of LifeAct::RFP for control and *arx-2(RNAi)* embryos at 180 s after NEBD. After *arx-2(RNAi)* F-actin puncta (white arrowheads) disappear.

**B)** Confocal single z-plane images of the cell cortex of NG::ANI-1 (green) and NMY-2::mKate (red) after *arx-2(RNAi)* or *cyk-1(RNAi); arx-2(RNAi)*. Magnification of the cell equator is shown on the right. Note: cleavage furrow ingression is delayed in *arx-2(RNAi)* in comparison to control embryos (compare with Fig. 2A, 180 s). Linear structures are highlighted by red arrowheads and non-linear by blue arrowheads.

**C)** Normalized mean NG::ANI-1 fluorescence intensity from the anterior to the posterior cortex at 180 s after NEBD for indicated RNAi conditions. Control graph is reproduced from Fig. S3B and n=number of embryos analyzed.

**D)** The mean number of linear and non-linear NG::ANI-1 structures per embryo at an equatorial region for the indicated RNAi conditions at 180 s after NEBD. P-values were calculated using Mann-Whitney test and represent ** p<0.01 in comparison to control embryos treated without RNAi, n≥5 embryos for each condition.

All scale bars are 5 μm and error bars are SEM.

**Figure S5 Single, double and triple RNAi co-depletions of NMY-2, CYK-1 and septin^UNC-61^ are highly efficient**

**A)** Confocal single z-plane cortical images of NMY-2::mKate for indicated RNAi conditions 180 s after NEBD.

**B)** Graphs display the mean normalized NMY-2::mKate cortical fluorescence intensity for the indicated proteins and RNAi conditions.

**C)** Immunoblots of adults worms expressing endogenously-tagged GFP::NMY-2 with or without *nmy-2(RNAi)* and probed with the indicated antibodies. The mean intensity of the GFP::NMY-2 band is shown (3 worm extracts).

**D)** Normalized cortical NG::ANI-1 fluorescence intensity for indicated RNAi conditions 180 s after NEBD. Control graph reproduced from Fig. S3B.

**E)** Central plane images of embryos expressing the general membrane marker mCherry::PH-PLCδ1 and mCherry::DNA (Histone-H2B) and treated with the indicated RNAi conditions. Error bars are SEM, time is indicated in seconds (s) after NEBD, scale bars are 5 μm and n= number of embryos analyzed.

**Figure S6 Cortical fluorescence intensities of NMY-2::mKate and NG::ANI-1 after Latrunculin A treatment**

**A-C)** Maximum intensity projections of 10 cortical z-planes of permeabilized one-cell embryos expressing NMY-2::mCherry (red) and LifeAct::GFP (green) treated with DMSO (A) or Latrunculin A (B, C). Panel (C) shows a kymograph of the equatorial region of the embryo in panel (B).

**D, E)** Maximum intensity projections of 10 cortical z-planes of permeabilized one-cell embryos expressing NMY-2::mKate (red) and NG::ANI-1 (green) treated with DMSO (D) or Latrunculin A (E). Panel (E) shows a kymograph of the equatorial region of the embryo of Fig. 4A.

**F)** Normalized cortical NMY-2::mKate and NG::ANI-1 fluorescence intensity from the anterior to the posterior cortex for control and Latrunculin A treated embryos at indicated time points. Error bars are SEM, scale bars are 5 μm and n=number of embryos analyzed.

**Figure S7 The N-terminal halves of *C. elegans* and human anillin are predicted to be highly disordered**

**A-B)** Graphs display predicted disordered regions of *C. elegans* ANI-1 and human anillin which were determine with DISOPRED3 (Jones and Cozzetto, 2015). For comparison a scheme of their functional domains is shown below.

**Figure S8 ANI-1^N-term^ and ANI-1^C-term^ mutant proteins partially rescue embryonic lethality after *ani-1(RNAi)***

**A)** Schematic representation of ANI-1^WT^ and tested ANI-1 mutant proteins.

**B)** GFP-tagged ANI-1 transgenes were integrated into the ‘Mos’ site on chromosome II. The ANI-1 transgenes comprise the genomic *ani-1* locus and *gfp*. Transgenes are resistant to *ani-1* RNAi targeting by re-encoding exon 5 but keeping the amino acid sequence and codon usage the same.

**C)** Graph is plotting the percentage of embryonic lethality for the indicated ANI-1 proteins and RNAi conditions. Error bars are SEM and n= number of progeny (larvae and embryos) counted.

**D)** Immunoblot of adult worms expressing the indicated ANI-1 proteins probed with anti-GFP and anti-actin antibodies.

**Figure S9 The Δ48-460 ANI-1 mutant forms linear structures after CYK-1 depletion**

**A-C)** Cortical single z-plane confocal images of the Δ48-160, Δ236-460 and Δ48-460 GFP-tagged ANI-1 mutants for the indicated RNAi conditions and time points after NEBD. Magnification of the cell equator is shown on the right. Linear structures are highlighted by red and non-linear by blue arrowheads. Since ANI-1 levels are reduced after septin^UNC-61^ co-depletion (Fig. 6B), intensity scaling was increased to better visualize the ANI-1 structures after *septin^unc-61^(RNAi)* (C). Scale bar is 5 μm.

**D)** Mean number of structures per embryo at the cell equator for the different GFP-tagged ANI-1 proteins after RNAi depletions 180 s after NEBD. P-values were calculated using Mann-Whitney or student t-test and represent ** p<0.01, *** p<0.001 in comparison to *ani-1(RNAi)* embryos expressing the same transgene, n≥5 embryos for each condition.

**E)** Mean percentage of 0-20° (anterior to posterior) and 68-88° (circumferentially) directionality measured for GFP-tagged ANI-1 proteins for indicated RNAi conditions 180 s after NEBD at the cell equator. *P*-values were determined with student *t*-test and represent ***p<0.001. ANI-1^WT^ graphs are reproduced from Fig. 5F.

All error bars are SEM.

**Figure S10 The Δ48-460 ANI-1 and ANI-1^C-term^ mutants do not support furrow ingression after co-depletion of septins and formin**

**A)** Merged central plane DIC and wide-field fluorescent images of the ANI-1^WT^, ANI-1^C-term^ and Δ48-460 ANI-1 proteins for the indicated RNAi conditions and time points after NEBD. Scale bar is 5 μm.

**B)** Plotted is the contractile ring diameter of each embryo over time for indicated RNAi conditions, n=number of embryos analyzed. Red encircled stars highlight that all embryos fail cytokinesis.

**Figure S11** Model illustrating how contraction of the ANI-1 network mediates cleavage furrow formation and ingression after the depletion of septins and formin.

## Movie Legends

**Movie 1, related to Fig. 1:** Central plane confocal images of one-cell *C. elegans* embryos expressing mCherry::PH-PLCδ1 and mCherry::DNA (Histone-H2B) markers treated with the indicated RNAi conditions. Images were acquired every 20 s on a Nikon eclipse Ti spinning disk confocal controlled by NIS Elements 4.51 software equipped with a 100x 1.45-NA Plan-Apochromat oil immersion objective and Andor DU-888 X11056 camera. Movie starts 140 s after NEBD.

**Movie 2+3, related to Fig. 2 and S4:** Confocal images of the cell cortex of one-cell *C. elegans* embryos expressing NG::ANI-1 (green) and NMY-2::mKate (red) treated without (control) or with *cyk-1(RNAi)* or with *cyk-1(RNAi); septin^unc-61^(RNAi)* (Movie 2), and with *arx-2(RNAi)* or with *cyk-1(RNAi); arx-2(RNAi)* (Movie 3). Note: Scaling intensity for NG::ANI-1 after septin^UNC-61^ co-depletion are increased. Images were acquired every 2.5 s on a Nikon eclipse Ti spinning disk confocal controlled by NIS Elements 4.51 software equipped with a 100x 1.45-NA Plan-Apochromat oil immersion objective and Andor DU-888 X11056 camera. Time in seconds (s) after NEBD is shown.

**Movie 4, related to Fig. 3:** Confocal images of the cell cortex of one-cell *C. elegans* embryos expressing NG::ANI-1 treated with the indicated RNAi conditions. Intensity scaling was increased to visualize the NG::ANI-1 structures after *cyk-1(RNAi); nmy-2(RNAi); septin^unc-61^ (RNAi)*. Imaging conditions are the same as indicated for Movie 2. Time in seconds (s) after NEBD is shown.

**Movie 5, related to Fig. 4 and S6:** Maximum z-projections of 10 confocal images of the cell cortex of one-cell *C. elegans* embryos expressing NG::ANI-1 (green) and NMY-2::mKate (red) treated without or with Latrunculin A. Images were acquired every 5 s on inverted microscope (Ti-E, Nikon) equipped with a 60x oil-immersion Plan-Apochromat objective (N.A. 1.4) and an iXon Ultra 897 camera. Time in seconds (s) after NEBD is shown.

**Movie 6, related to Fig. 4:** Maximum z-projections of 10 confocal images of the cell cortex of one-cell *C. elegans* embryo expressing GFP::ANI-1^AH-PH^ (green) and NMY-2::mKate2 (red) treated with Latrunculin A and *ani-1(RNAi)*. Images were acquired as described for Movie 5. Time in seconds (s) after NEBD is shown.

**Movie 7, related to Fig. 5:** Confocal images of the cell cortex of one-cell *C. elegans* embryos expressing indicated GFP::ANI-1 variants treated with *ani-1(RNAi)*. Images were acquired every 5 s on laser scanning microscope (SP8) equipped with a 63x/1.3 GLYC objective and photomultiplier. Time in seconds (s) after NEBD is shown.

**Movie 8, related to Fig. 5:** Confocal images of the cell cortex of one-cell *C. elegans* embryos expressing indicated GFP::ANI-1 variants treated with *cyk-1(RNAi) ani-1(RNAi)*. Note: Fluorescence intensity scalings are increased for ANI-1^N-term^ in comparison to ANI-1^WT^. Images were acquired as described for Movie 7. Time in seconds (s) after NEBD is shown.

**Movie 9, related to Fig. 5:** Confocal images of the cell cortex of one-cell *C. elegans* embryos expressing indicated GFP::ANI-1 variants treated with *cyk-1(RNAi) ani-1(RNAi); septin^unc-61^ (RNAi)*. Images were acquired as described for Movie 7. Time in seconds (s) after NEBD is shown.

**Movie 10-12, related to Fig. S9:** Confocal images of the cell cortex of one-cell *C. elegans* embryos expressing indicated GFP::ANI-1 variants treated with *ani-1(RNAi)* (Movie 10), *cyk-1(RNAi) ani-1(RNAi)* (Movie 11) or *cyk-1(RNAi) ani-1(RNAi); septin^unc-61^ (RNAi)* (Movie 12). Images were acquired as described for Movie 7. Time in seconds (s) after NEBD is shown.

## Material and Methods

### *C. elegans* maintenance and RNAi experiments

*C. elegans* strains were maintained at 20°C on NGM plates seeded with *Escherichia coli* (OP50) (Stiernagle, 2006). A summary of the worm strains used is provided in Table S1.

RNAi experiments were performed by injection or feeding and details of the RNAi experiments are listed in Table S2. For the generation of dsRNA for injection the target sequence was amplified by PCR from cDNA or genomic DNA with sequence specific primers containing T7 overhangs on both sides (Table S2). The purified PCR product was used as a template for the *in vitro* transcription reaction using the MEGAscript T7 kit (AM1334; Invitrogen). dsRNA was injected into young hermaphrodites mounted on 2 % agarose pads. After injections worms were rescued on OP50 seeded NGM plates and incubated at 20°C for 24-28 h *(nmy-2)*, or 39-49 h *(cyk-1, ani-1, arx-2, septin^unc-59^, septin^unc-61^*).

For feeding RNAi, L4440 vectors carrying part of the sequence of *ani-1* or *perm-1* were obtained from the Ahringer library (Source Bioscience), sequenced to confirm the gene target, and transformed into HT115 bacteria. Briefly, the RNAi bacterial clones were grown until an OD of 1.6 in 50 ml Lysogeny broth (LB) medium containing 12.5 μg/ml tetracycline and 50 μg/ml ampicillin overnight at 37°C. The overnight culture was centrifuged at 4000 g for 10 min, the supernatant was removed, and the cell pellet was resuspended in 2.5 ml LB medium containing 12.5 μg/ml tetracycline, 50 μg/ml ampicillin, and 0.23 mg/ml IPTG (isopropyl β-D-1-thiogalactopyranoside). In parallel, unseeded NGM plates were dried for 1 h in a 37°C incubator. Then, 100 μl of a 1:1:1 mix of 50 μg/ml ampicillin, 12.5 μg/ml tetracycline, and 0.1 μg/ml IPTG was added. These treated plates were inoculated with 100 μl of bacterial culture. To obtain permeabilized embryos in Fig. 4, and S6, the inoculated bacterial culture consisted of 20 μl of bacteria expressing *perm-1* dsRNA (Carvalho et al., 2011) and 80 μl of LB *(perm-1(RNAi)* alone), or 20 μl of bacteria expressing *perm-1* dsRNA and 80 μl of bacteria expressing *ani-1 (perm-1(RNAi))*. The expression of dsRNA was induced for 5 h at 37°C, in the dark. 25–30 L4 stage hermaphrodites were added to the treated plates and incubated at 20°C for 45-48 h before dissection for imaging.

### Embryonic lethality counts

To determine embryonic lethality after *ani-1(RNAi)*, young adult worms were injected with *ani-1* dsRNA and incubated at 20°C for 48 h. Afterwards, injected worms were singled on NGM plates, incubated 20 h and sacrificed after 24 h. The number of larvae and dead embryos was counted 24 h later.

### Immunoblotting

For immunoblotting, worms were picked in MPEG (M9 plus 0.05% polyethylene glycol 8000 (PEG)) buffer and washed three times. The volume of the MPEG buffer was reduced and sample buffer was added to reach approximately 1 worm/μl. Worms were incubated at 95°C for 5 min, followed by centrifugation at room temperature for 10 min at 20000 *g* to spin down debris and subsequently 20 μl of supernatant were loaded in each lane (Zanin et al., 2011). Membranes were incubated with anti-Actin (A-1978, Sigma, 1:6000), anti-FLAG (F1804 Sigma; 1:2500) or anti-GFP (11814460001 Roche; 1:600) primary antibodies. HRP-conjugated rabbit (170-6515 Bio-Rad; 1:15000) and HRP-conjugated mouse (170-6516 Bio-Rad; 1:7500) were used as secondary antibodies.

### Mos1-mediated single copy insertion (MosSCI) of ANI-1 variants

For the *gfp::ani-1* transgenes the genomic *ani-1* locus (6507 bp) with its promotor (565 bp) and 3’UTR (319 bp) were cloned together with GFP into pCFJ350 by Gibson cloning. Exon 5 was rendered RNAi resistant by amino acid codon shuffling (Fig. S8B). For the ANI-1^N-term^ variant the PBS and the CX motif of RHO-1 (AA181-192) were fused to AA764 of ANI-1. We chose this region as a membrane anchor, since it was sufficient for membrane localization and did not exhibit any specific cortical enrichment during anaphase (not shown). All transgenes are listed in Table S3.

*Ani-1* transgenes were integrated on chromosome II (EG8079) using the MosSCI method (Frøkjær-Jensen et al., 2008). Young hermaphrodites were mounted on 2 % agarose pads and injected with a mix containing the vector carrying the *ani-1* transgene, the transposase (pCFJ601) and co-injection markers (pCFJ90 and pCFJ104). Worms were singled on NGM plates seeded with OP50 and incubated at 25°C for 7-10 days until starved. Wild type movers, negative for the mCherry-tagged array markers, were checked for homozygous GFP expression and correct integration was verified using PCR.

### Fluorescence microscopy and image analysis

*C. elegans* gravid hermaphrodites were dissected in 4 μl M9 buffer on a 18×18 mm coverslip. The embryos were mounted on a 2 % agarose pad by inverting the coverslip onto the pad and then sealed with vaseline. Confocal images (Fig. 1–3, S1-S5) were acquired using a Nikon inverted microscope (Eclipse Ti) equipped with a confocal spinning disk unit, a 100x 1.45-NA Plan-Apochromat oil immersion objective and an Andor DU-888 X-11056 camera (1024×1024 pixels). The system was controlled by NIS Elements software. GFP and red-fluorescent probes were imaged using 488 nm and 561 nm lasers, respectively, and for some experiments transmission images were acquired in the central plane. Confocal images of Fig. 5, S9 were acquired on the Leica SP8 confocal laser scanning microscope equipped with a white light laser, 63x/1.3 GLYC objective and a photomultiplier. To display images acquired on the SP8, a Gaussian Blur filter with a radius of 0.8 was applied to reduce noise. Cortical images in Fig. 4, S6 were acquired on a spinning disk confocal system (Andor Revolution XD Confocal System; Andor Technology) with a confocal scanner unit (CSU-X1; Yokogawa Electric Corporation) mounted on an inverted microscope (Ti-E, Nikon) equipped with a 60x 1.4 NA Plan-Apochromat oil objective and solid-state lasers of 488 nm (50 mW) and 561 nm (50 mW). For image acquisition, 12×0.5 μm z-stacks were collected every 5 s by using an electron multiplication back-thinned charge coupled device camera (iXon Ultra 897; Andor Technology). Acquisition parameters, shutters and focus were controlled by Andor iQ3 software. Wide-field fluorescent and DIC images in Fig. 6 and S10 were acquired on the Nikon Ti2-Eclipse microscope equipped with a CFI Apochromat 100x 1.49 NA oil objective and a Prime 95B, A18E203001 camera.

Image analysis and quantification was performed in Fiji (Schindelin et al., 2012). All data analysis was performed in Excel, Prism (GraphPad) or KNIME analytics (http://www.knime.org) and figures were assembled with the Affinity Designer software.

For high-time resolution data, imaging was started during pronuclear migration with central planes images acquired every 16 s (Fig. 2–3, 5, S1, S2D, S3-4, S5A, S9). At the time of NEBD (defined as the time point when the border of the nucleus was no longer visible in the DIC channel) imaging was stopped and resumed at the cell cortex before anaphase onset with a time interval of 2.5 s (Fig. 2–3, S1, S2D, S3-4, S5A) or 5 s (Fig. 5, S9).

Fluorescence intensity along cortical plane images (Figs. 5B, S1B, D, S2E, S3B, D, S4C, S5B, S6F,) was measured on raw images by drawing a wide line from the anterior to the posterior cortex of the embryo. Cortical fluorescence intensity was normalized by dividing by the mean cytoplasmic intensity measured in a rectangular box in the posterior region of central plane images at the time of NEBD. The mean normalized cortical intensity was calculated for 101 intervals from the anterior (0%) to the posterior (100%) for each embryo. To correct the fluorescent signal for bleaching, the mean cortical intensity was measured over time in a rectangular box in the posterior region for each embryo. The mean fluorescence intensity relative to the initial intensity was calculated for control embryos (n≥11) for each time point and used as a ‘correction coefficient’. To correct for bleaching, the fluorescent intensity measured at a specific time point was multiplied with the ‘correction coefficient’ of this time point.

Directionality of ANI-1 and LifeAct was determined on cortical images that were processed by subtracting the mean fluorescence intensity measured outside the embryos. Subsequently a Gaussian Blur filter with a radius of 0.5 was applied followed by an Unsharp Mask filter with a radius of 2 pixels and a Mask Weight of 0.6. Images were also rotated (anterior – left, posterior – right). To measure directionality a rectangular box was placed at the cell equator at selected time points after NEBD and the ImageJ plugin ‘Directionality’ with the method local gradient orientation (Liu, 1991) was used. The orientation of ANI-1 and LifeAct was classified into 45 bins from 0–90° orientation. The mean percentage of ANI-1 and LifeAct structures with 0-20°, reflecting anterior-posterior orientation, and 68-88°, reflecting circumferentially orientation, was calculated for each embryo.

The number of linear and non-linear structures was analyzed on cortical images that were processed as described for the directionality analysis. Additionally, the mean fluorescence intensity measured at the anterior cortex was subtracted from the cortical image at the end. A rectangular box was positioned at the cell equator (anterior for ANI-1^N-term^ embryos) and the length and width of all structures was determined manually. For each structure the length/width ratio was calculated and structures with the length/width ratio <4 were classified as non-linear and structures with a length/width ratio ≥4 were classified as linear.

The contractile ring diameter was measured manually on central plane images acquired every 20 s starting at NEBD. The initial ring diameter NEBD was set to 100% diameter width. Embryos were filmed from NEBD of the first cell division until to onset of the second cell division.

### Latrunculin A treatment

Adult hermaphrodites that had been fed bacteria expressing *perm-1* dsRNA (see above) were dissected, and permeabilized embryos were filmed in meiosis medium (25 mM HEPES, pH 7.4, 0.5 mg/ml inulin, 20% heat-inactivated fetal bovine serum, and 60% Leibowitz-15 medium) without compression (Carvalho *et al*., 2011). 10 μM Latrunculin A (Sigma) was added between metaphase and anaphase onset. At the end of each video, medium containing 33 μM FM4-64 (Molecular Probes) was added to the imaging chamber to confirm that the imaged embryo was permeable.

### 6xHIS::NMY-2 cloning and NMY-2/MLC-4/MLC-5 protein expression

NMY-2 full length was amplified from *C. elegans* cDNA and cloned into the pACEbac1 expression vector using Gibson assembly with an N-terminal 6×Histidine tag followed by a flexible linker (GlySerGlySerGly). The primers used were oAC1233 - 5’-ACCATGGCTCTGGTAGCGGCACATCATCTCGACAAAAAGATGATGAG-3’ (forward) and oAC1234 - 5’-TAGTACTTCTCGACAAGCTTTTAGTTGCGAACTGAGTCGCG GTCT-3’ (reverse). The backbone was amplified from pACEbac1 vector using the following primers oAC1235 – 5’-AAGCTTGTCGAGAAGTACTAGAGGATCATAATC-3’ (forward) and oAC1236 5’-GCCGCTACCAGAGCCATGG-3’ (Reverse). Baculovirus expression of the 6xHIS::NMY-2 full-length / StreptagII::MLC-4 / StreptagII:MLC-5 complex, cell lysates and tandem streptactin/Ni-NTA affinity chromatography were performed as described previously for NMY-2 HMM complexes (Osório *et al*., 2019). The concentrated eluate of the Ni-NTA purification was dialysed overnight in high salt buffer (10 mM MOPS, 500 mM NaCl, 3 mM NaN3, 1 mM EDTA, 1 mM DTT (pH 7.3)). Glycerol and DTT were added to a final concentration of 10% (vol/vol) and 1 mM, respectively. Aliquots were flash-frozen in liquid nitrogen and stored at −80°C.

### PIP strips and immunoblotting

To analyze lipid binding, PIP strips (Echelon Bioscience, P-6001) were used according to the manufacturer’s instructions. Each membrane consists of 16 spots: 15 contain several lipids and 1 empty (blank). Additionally, on the empty space on the top of the membranes, we spotted two controls for proper detection consisting of 1 μL of each protein sample (0.5 – 1 μg) or 1 μL of the secondary antibody pre-diluted 1:100 (secondary antibody control). The two control spots were allowed to dry before proceeding with the protocol. Membranes were then incubated with blocking solution (3% BSA fatty-acid free (Sigma-Aldrich) in PBS (Gibco, Thermo-Fisher) with 0.1% (v/v) Tween 20) (PBS-T) for one 1 hr at RT. Membranes were then incubated with 4 μg/mL of either StreptagII::MLC-4 or 6xHIS::NMY-2/ Strep-tag II::MLC-4/ Strep-tag II:MLC-5 complex in blocking buffer for 1 h at RT, followed by three washes in PBS-T with gentle agitation for five minutes. To detect bound proteins, membranes were incubated for 1 h at room temperature in blocking solution with mouse StrepMAB antibody (IBA Lifesciences, 1:750, 2-1507-001) for Strep-tagII::MLC-4 detection or mouse anti-6xHistidine tag antibody (Millipore clone HIS.H8, 1:2,500, 05-949) for 6xHIS::NMY-2 detection. After washing, membranes were incubated with HRP-conjugated anti-mouse antibody at 1:5,000 in blocking solution and incubated for 1 h at RT. This was followed by another round of three washes and PBS was used for the final wash. All steps were performed under gentle agitation. Immunoblots were visualized by chemiluminescence using Pierce ECL Western Blotting Substrate (Thermo Fisher Scientific) and a ChemiDoc XRS+ System with Image Lab Software (Bio-Rad).

### Statistical analysis

Statistical analysis was performed in Prism (GraphPad). Mean values are presented with error bars representing standard error of the mean (SEM) as indicated in the figure legends. For data with normal distribution a parametric (two sided Student’s *t*-test) and with non-normal distribution a Mann-Whitney test for was performed.

---

**Table S1.** *C. elegans* strains used in the study.

| Strain name | Genotype | Source |
| --- | --- | --- |
| N2 | Wild type (ancestral) |  |
| OD70 | *Itls44[pie-1p::mCherry::PH(PLC1delta1)+unc-119(+)]* | (Kachur et al., 2008) |
| MG617 | *xsSi5 [pie-1p::GFP::ani-1<sup>AH-PH</sup>::pie-1 3'UTR + Cbr-unc-119(+)]II* | (Tse et al., 2012) |
| MDX29 | *ani-1(mon7[mNeonGreen<sup>3xFlag</sup>::ani-1]) III* | (Rehain-Bell et al., 2017) |
| BV70 | *zbls2[pie-1p::LifeAct::RFP + unc-119(+)]* | Zhirong Bao |
| LP229 | *nmy-2(cp52[nmy-2::mkate2 + LoxP unc-119(+)] LoxP) I; unc-119(ed3) III* | (Dickinson et al., 2017) |
| SWG004 | *cyk-1::gfp* | (Reymann et al., 2016) |
| EG8079 | *oxTi179 II; unc-119(ed3) III* | (Frøkjær-Jensen et al., 2008) |
| NK2225 | *unc-59(qy50[unc-59::GFP::3xflag::AID]) I* | (Chen et al., 2019) |
| ZAN23 | *nmy-2(cp13[nmy-2::gfp + LoxP] I; OD56 (mCherry::histone H2B; Itls44[pie-1p::mCherry::PH(PLC1delta1)+unc-119(+)]* | This study |
| ZAN348 | *ani-1(mon7[mNeonGreen<sup>3xFlag</sup>::ani-1]) III; zbls[pie-1::LifeAct::RFP + unc-119(+)]* | This study |
| ZAN351 | *nmy-2(cp52[nmy-2::mkate2 + LoxP unc-119(+)] LoxP) I; unc-119(ed3) III; ani-1(mon7[mNeonGreen<sup>3xFlag</sup>::ani-1]) III* | This study |
| ZAN368 | *nmy-2(cp52[nmy-2::mkate2 + LoxP unc-119(+)] LoxP) I; unc-119(ed3) III; xsSi5 [pie-1p::GFP::ani-1<sup>AH-PH</sup>::pie-1 3'UTR + Cbr-unc-119(+)] II* | This study |
| ZAN371 | *cyk-1::gfp, zbls2[pie-1p::LifeAct::RFP + unc-119(+)]* | This study |
| ZAN373 | *Si192[pEZ379; pani-1::GFP:ANI-1<sup>RE</sup>-WT; cb-unc-119(+)]II; unc-119(ed3) III* | This study |
| ZAN375 | *Si193[pEZ388; pani-1::GFP:ANI-1<sup>RE</sup>-Δ236-460AA; cb-unc-119(+)]II; unc-119(ed3) III* | This study |
| ZAN376 | *Si194[pEZ389; pani-1::GFP:ANI-1<sup>RE</sup>-Δ48-460AA; cb-unc-119(+)]II; unc-119(ed3) III* | This study |
| ZAN379 | *Si197[pEZ390; pani-1::GFP:ANI-1<sup>N-term</sup> (1-764AA-PBS-CX); cb-unc-119(+)]II; unc-119(ed3) III* | This study |
| ZAN382 | *Si200[pEZ391; pani-1::GFP:ANI-1<sup>C-term</sup> (Δ48-680AA); cb-unc-119(+)]II; unc-119(ed3) III* | This study |
| ZAN383 | *Si201[pEZ387; pani-1::GFP:ANI-1<sup>RE</sup>-Δ48-160; cb-unc-119(+)]II; unc-119(ed3) III* | This study |
| GCP22 | *Itls157 [pAC16;Ppie-1::LifeAct::GFP; unc-119 (+)]; unc-119(ed3) III; prtSi2[pAC71; Pnmy-2:: re-encoded nmy-2::mCherry::StrepTagII::3'UTRnmy-2; cb-unc-119(+)] II* | (Osório *et al.*, 2019) |
| GCP1126 | *unc-59(qy50[unc-59::GFP::AID])I; Itls37 [pAA64; pie-1/mCherry::his-58; unc-119 (+)]* | This study |

**Table S2.**
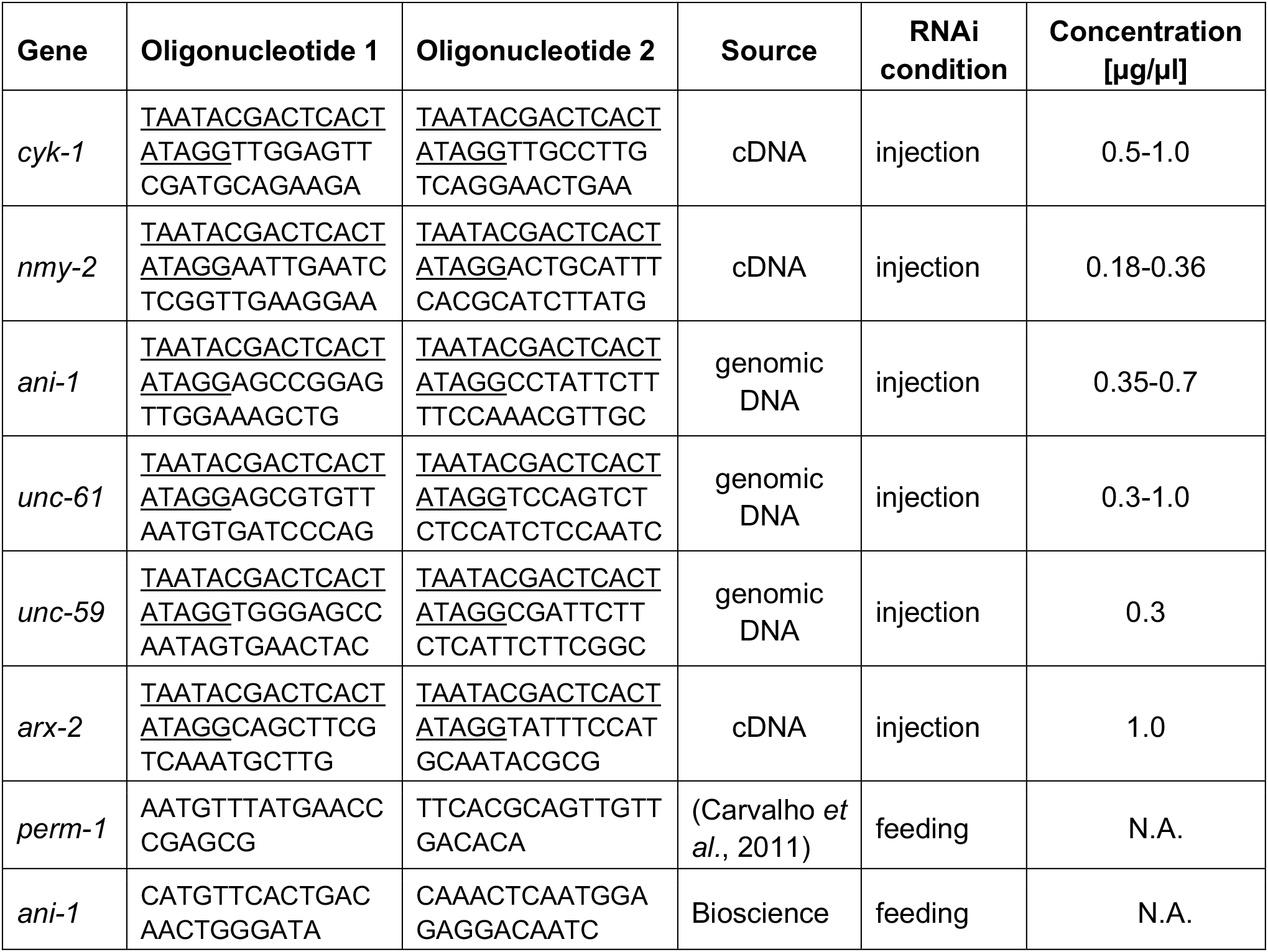
**dsRNA generation**, T7 sequence is underlined.

**Table S3.**
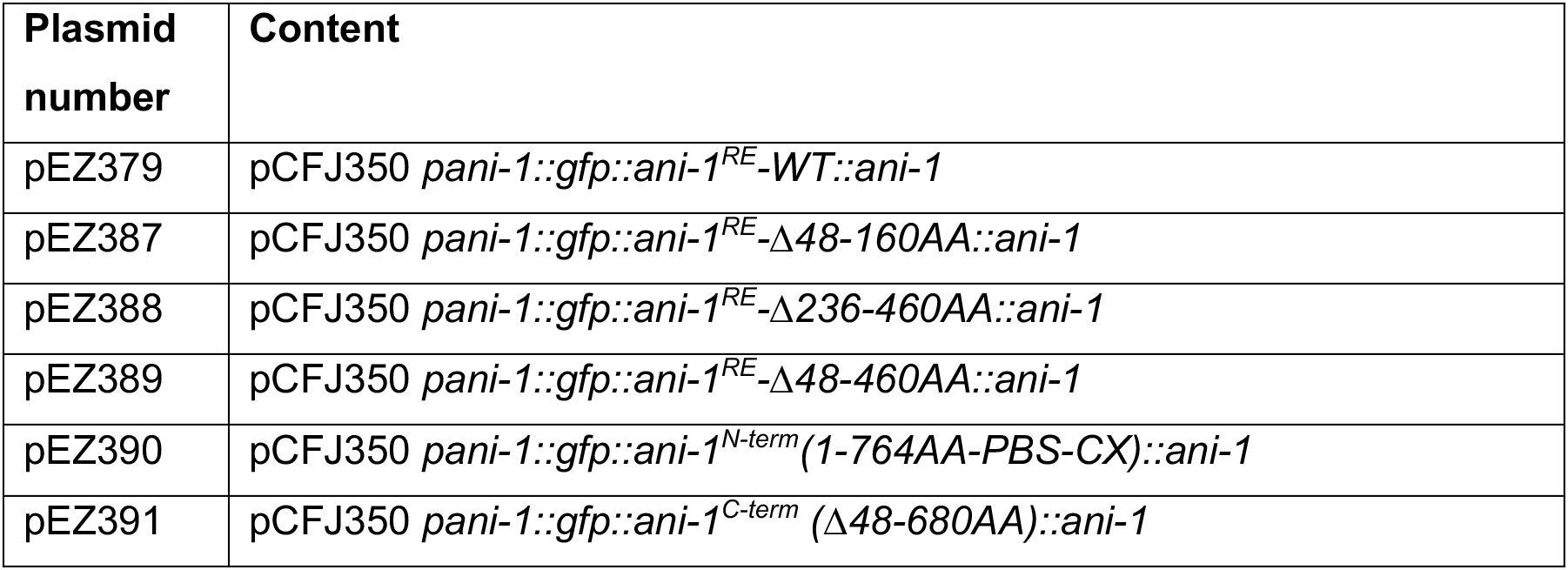
Generated Recombinant DNA.

## Acknowledgments

We are grateful to Christin Nöcker for technical assistance and to Maysoon Noureddine, Suraj Shaji, and Isabel Schultz-Pernice for helping with experiments. Microscopy was performed at the center for advanced light microscopy (CALM) at the LMU and for help with microscopy we thank Hartmann Harz and Christophe Jung. We are grateful to Christof Osman to access the Nikon Ti2-Eclipse microscope. For critical comments on the manuscript we thank Friederike Wolff and Sabine Müller. The Deutsche Forschungsgemeinschaft supported E. Zanin (ZA619/3), and T. Mikeladze-Dvail (MI1867/1-3). Research in AXC lab was funded by the European Research Council under the European Union’s Horizon 2020 Research and Innovation Programme (grant agreement 640553 – ACTOMYO) and AXC is supported by a Principal Investigator position from the Portuguese Foundation of Science and Technology FCT (CEECIND/01967/2017). F-Y. Chan has an FCT junior researcher position (DL 57/2016/CP1355/CT0013). J. Bellessem and E. Rackles were members of the Life Science Munich graduate program. For sharing *C. elegans* strains we thank Amy Maddox, Stephan Grill and Zhirong Bao. Some strains were provided by the CGC, which is funded by NIH Office of Research Infrastructure Programs (P40 OD010440).

