## Supplemental Figure 1-11 for "Anillin forms linear structures and facilitates furrow ingression after septin and formin depletion"

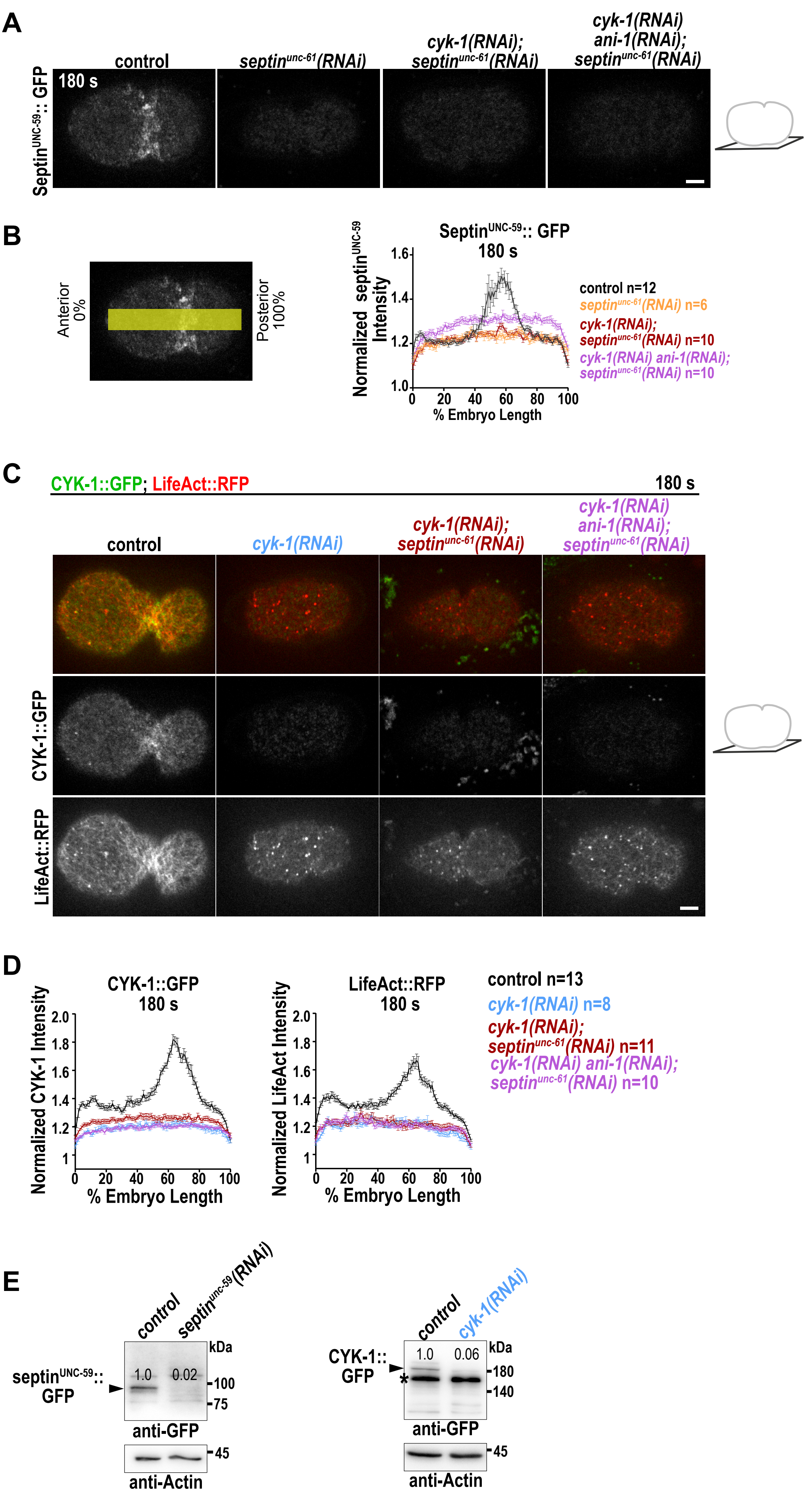

#### Supplemental FIGURE 2

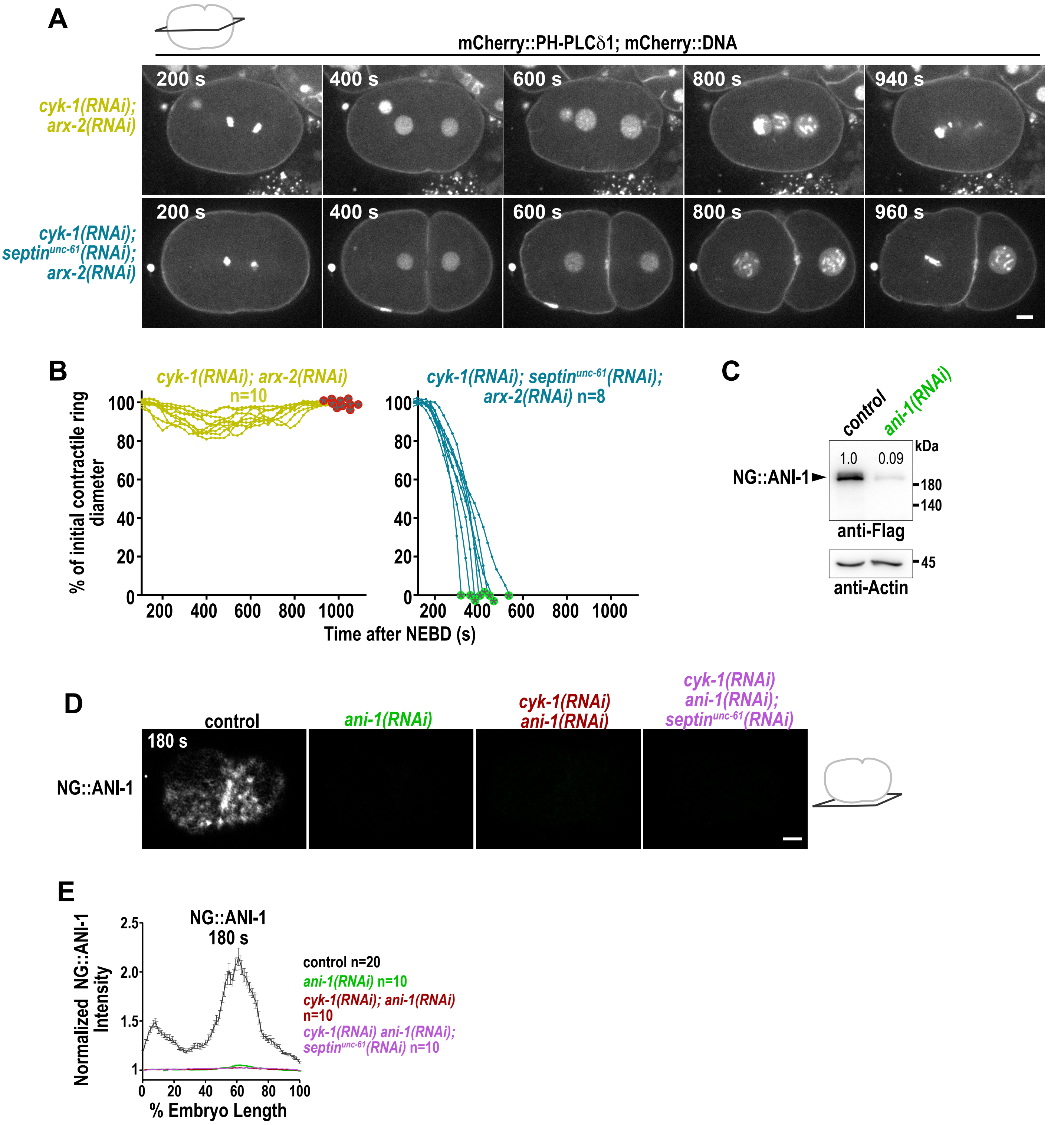

### Supplemental FIGURE 3

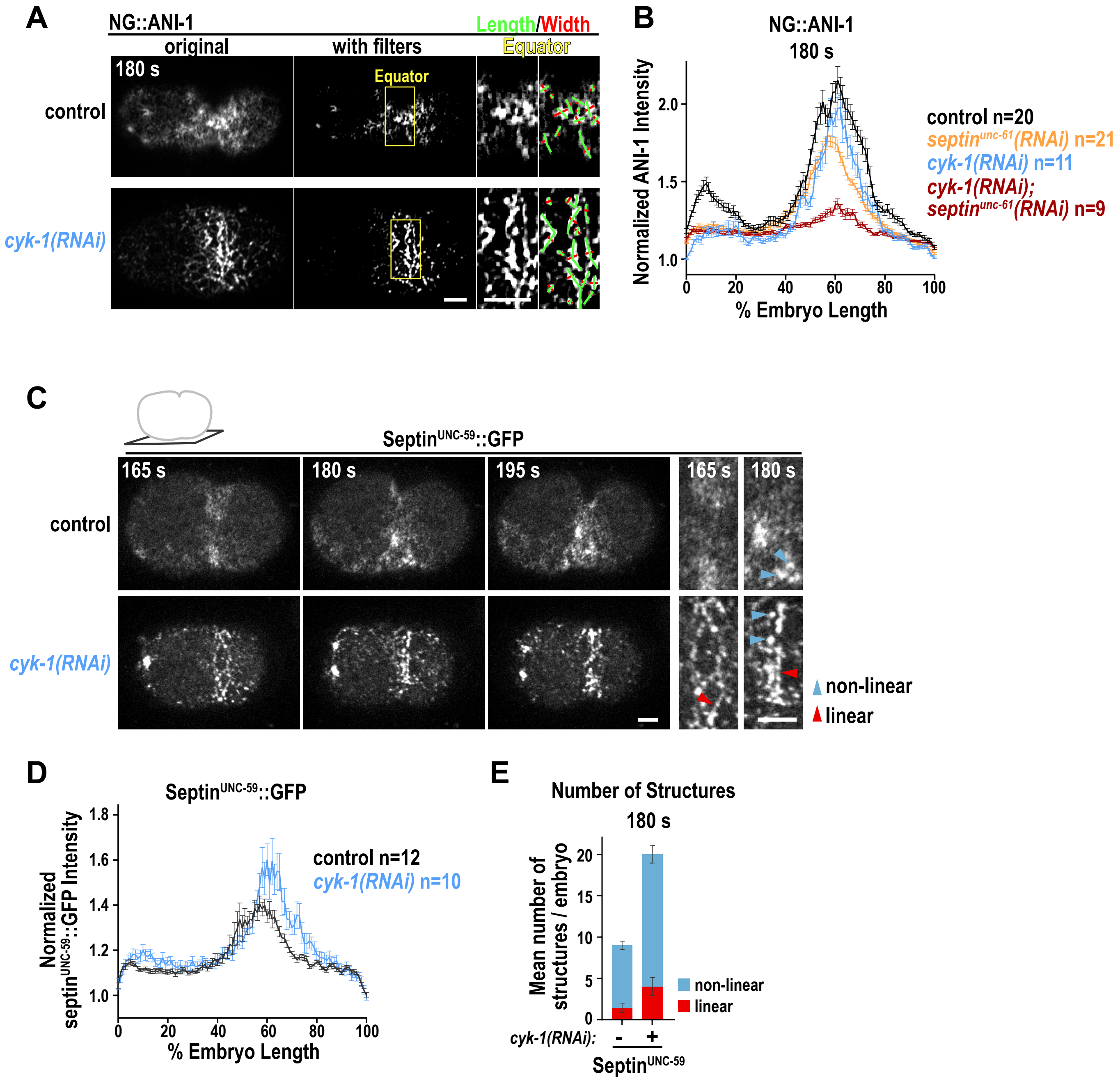

#### Supplemental FIGURE 4

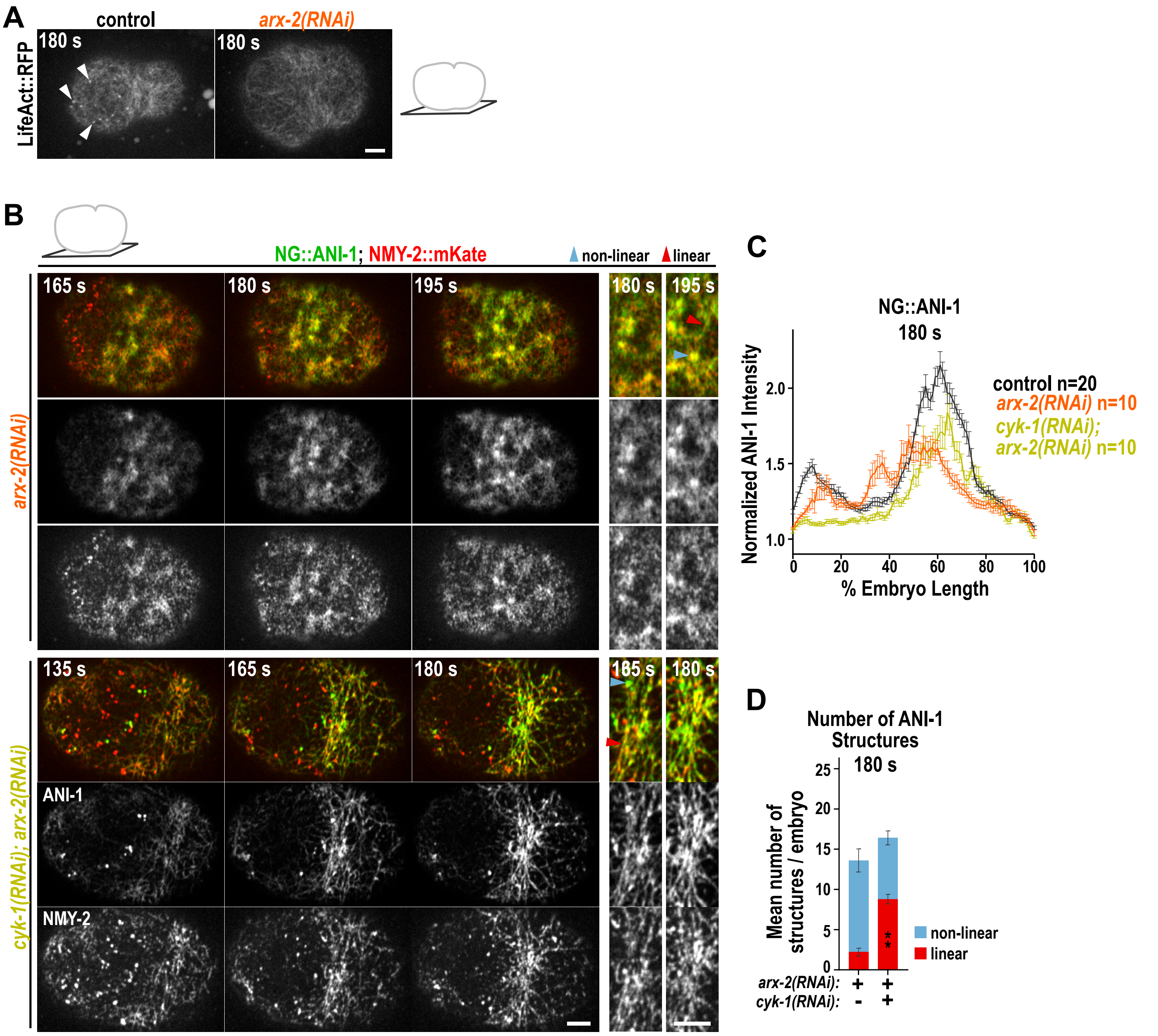

### Supplemental FIGURE 5

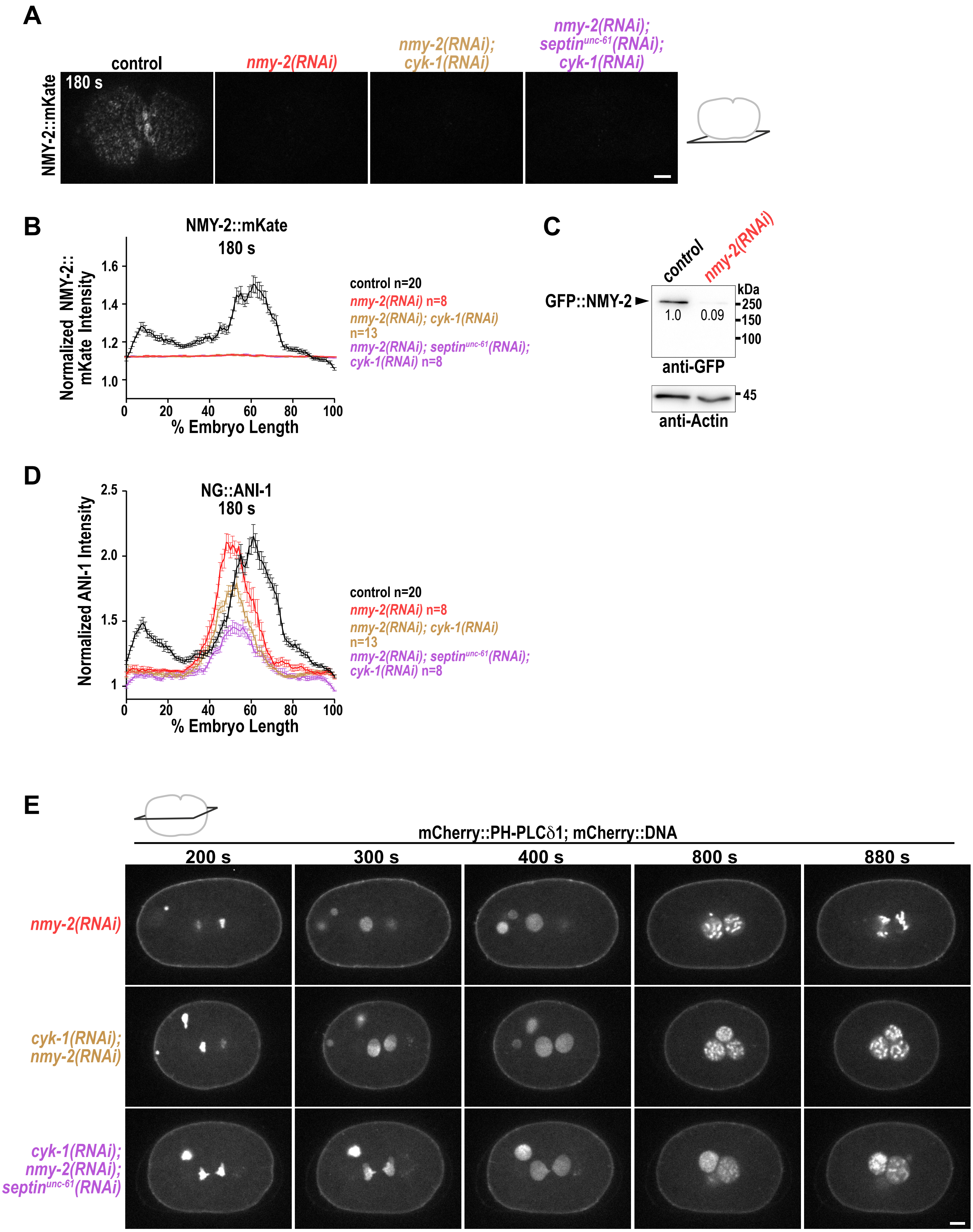

### Supplemental FIGURE 6

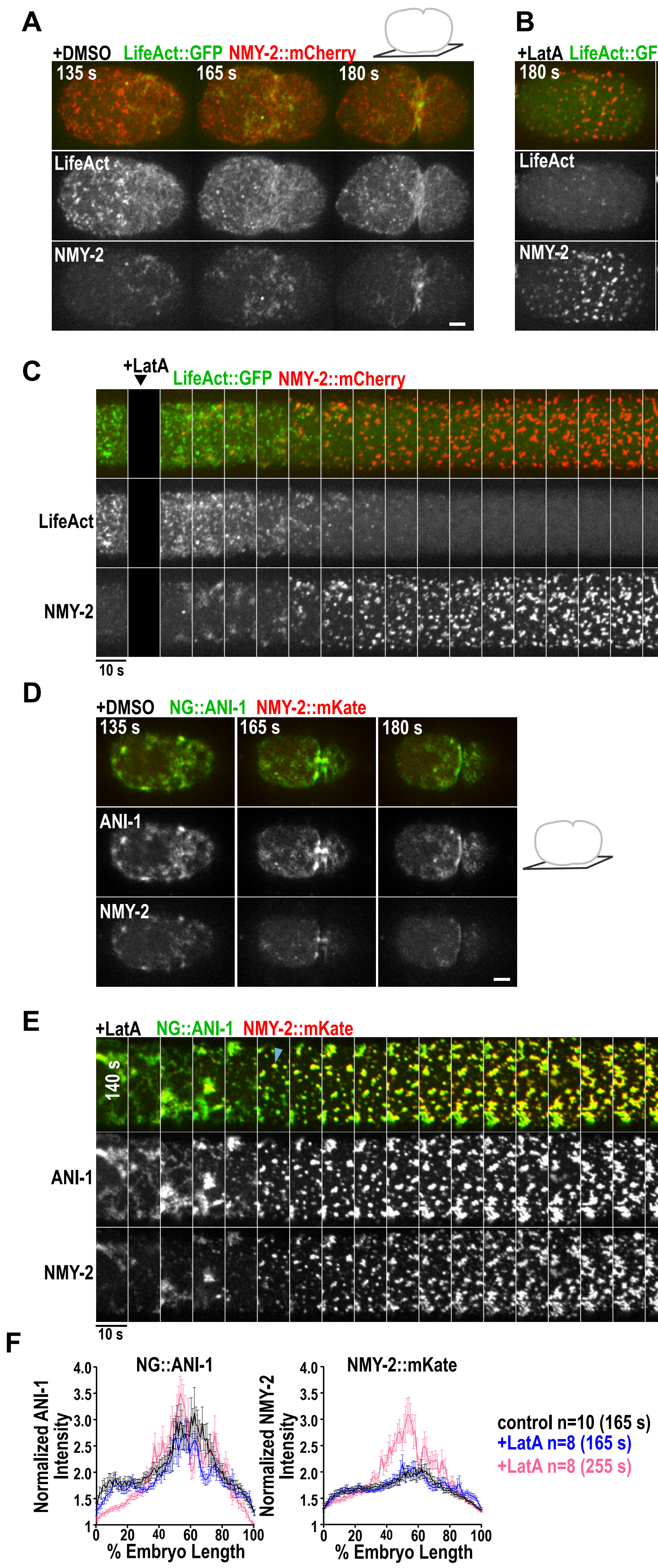

### Supplemental FIGURE 7

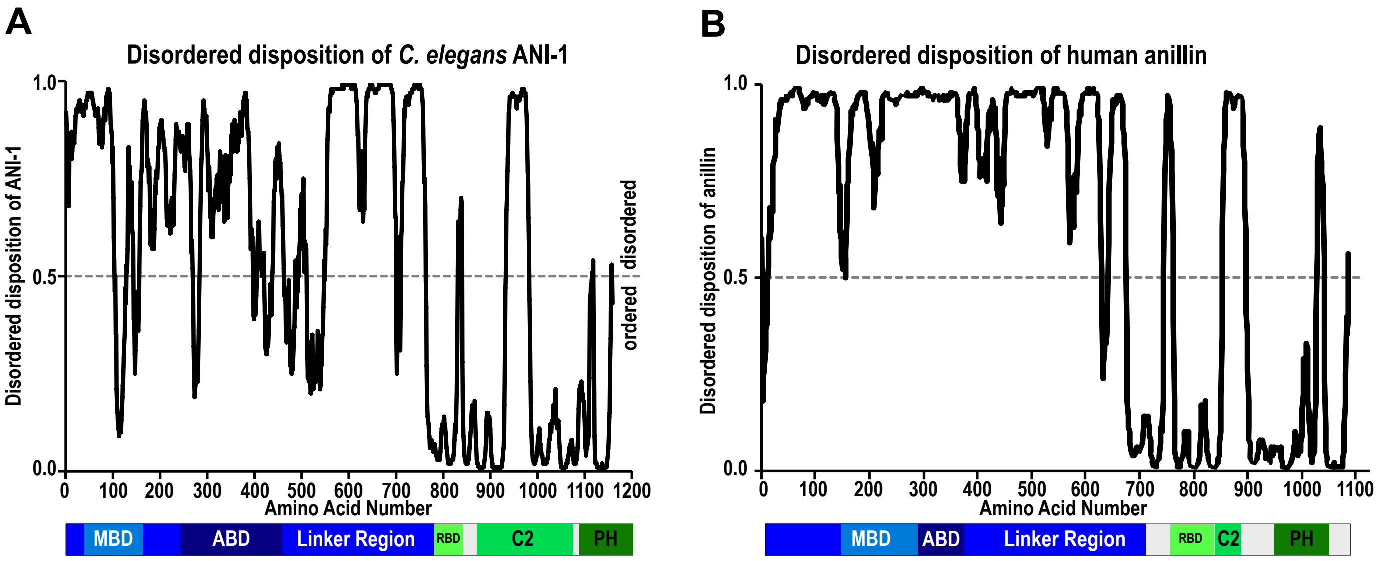

### Supplemental FIGURE 8

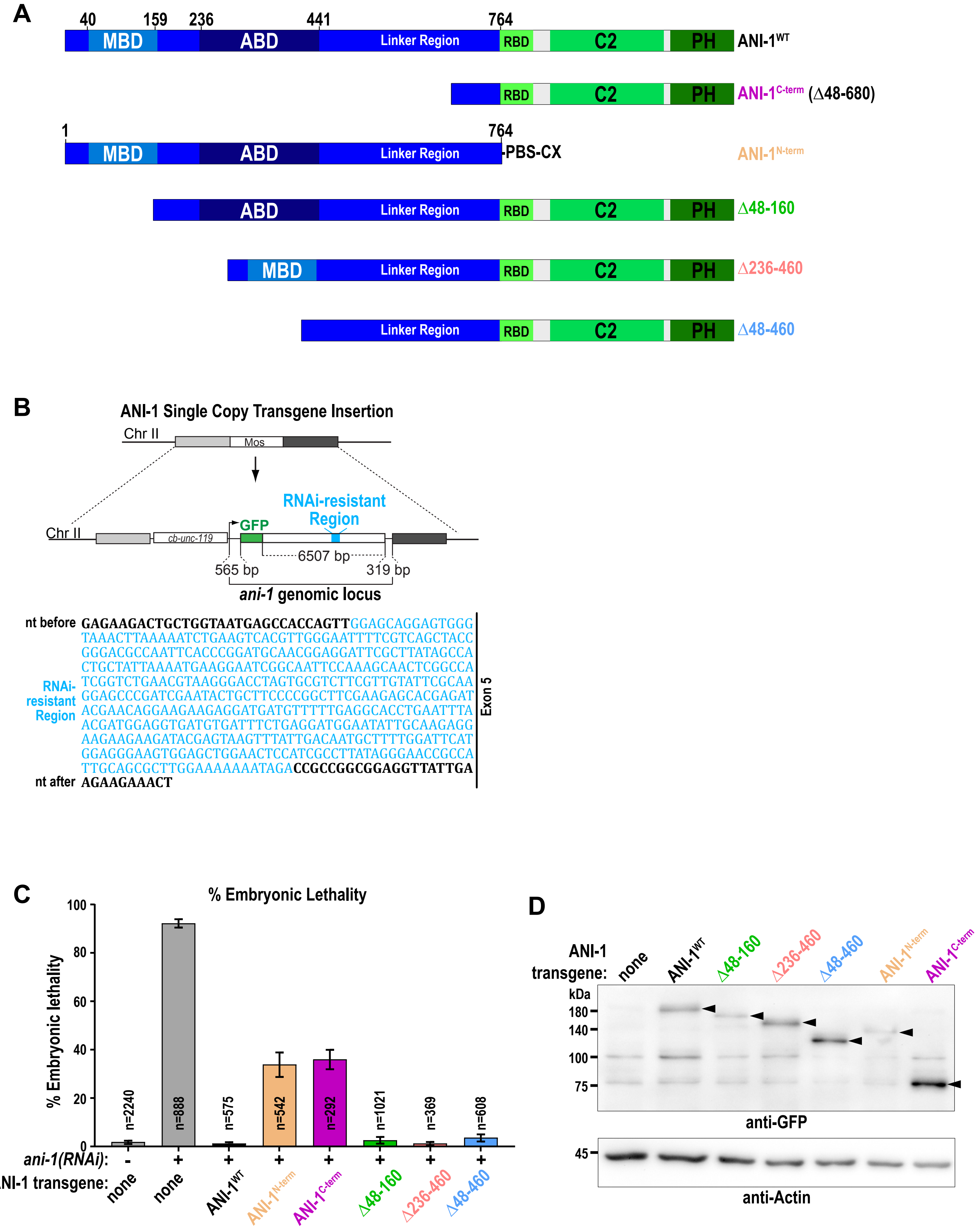

Supplemental FIGURE 9

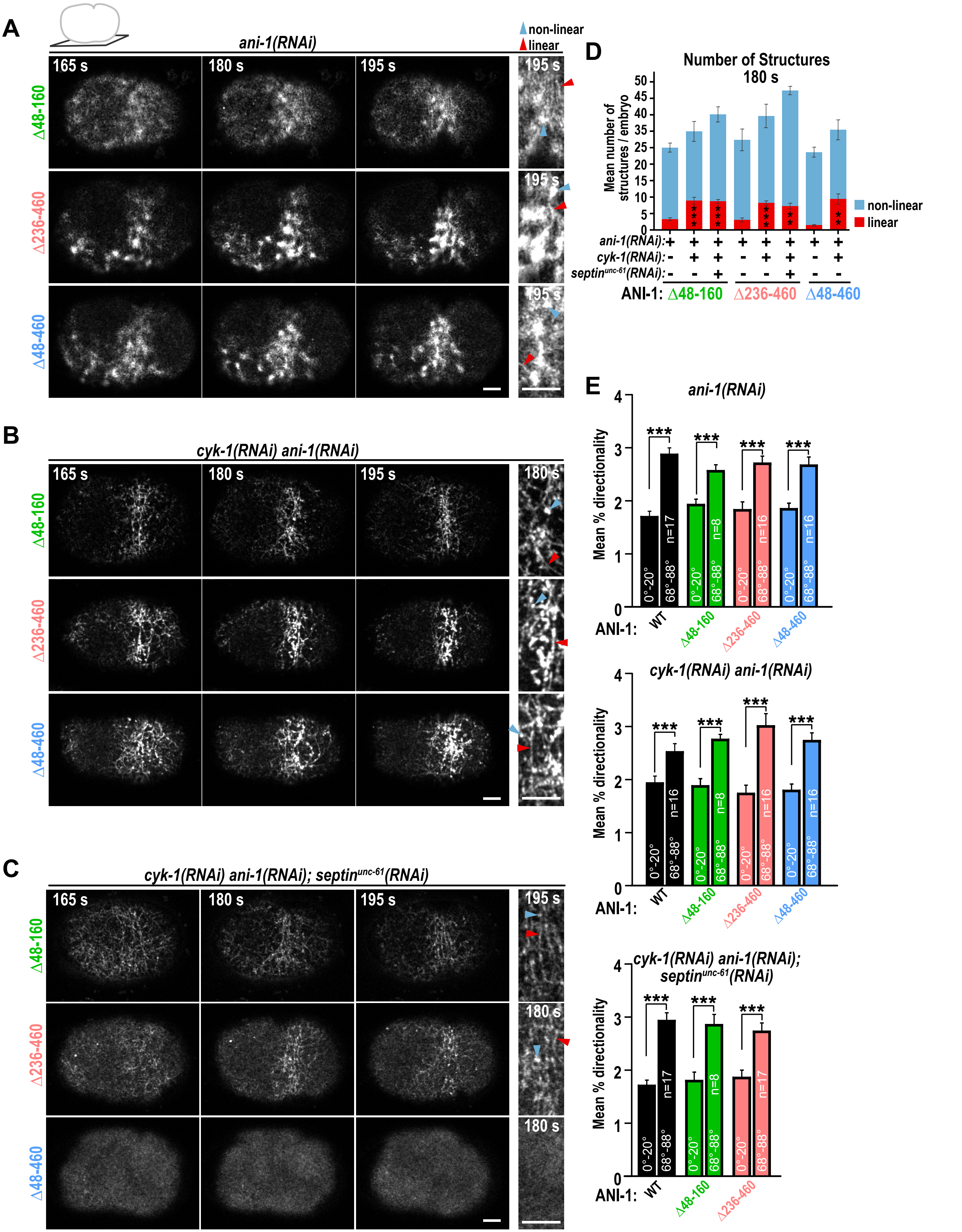

### Supplemental FIGURE 10

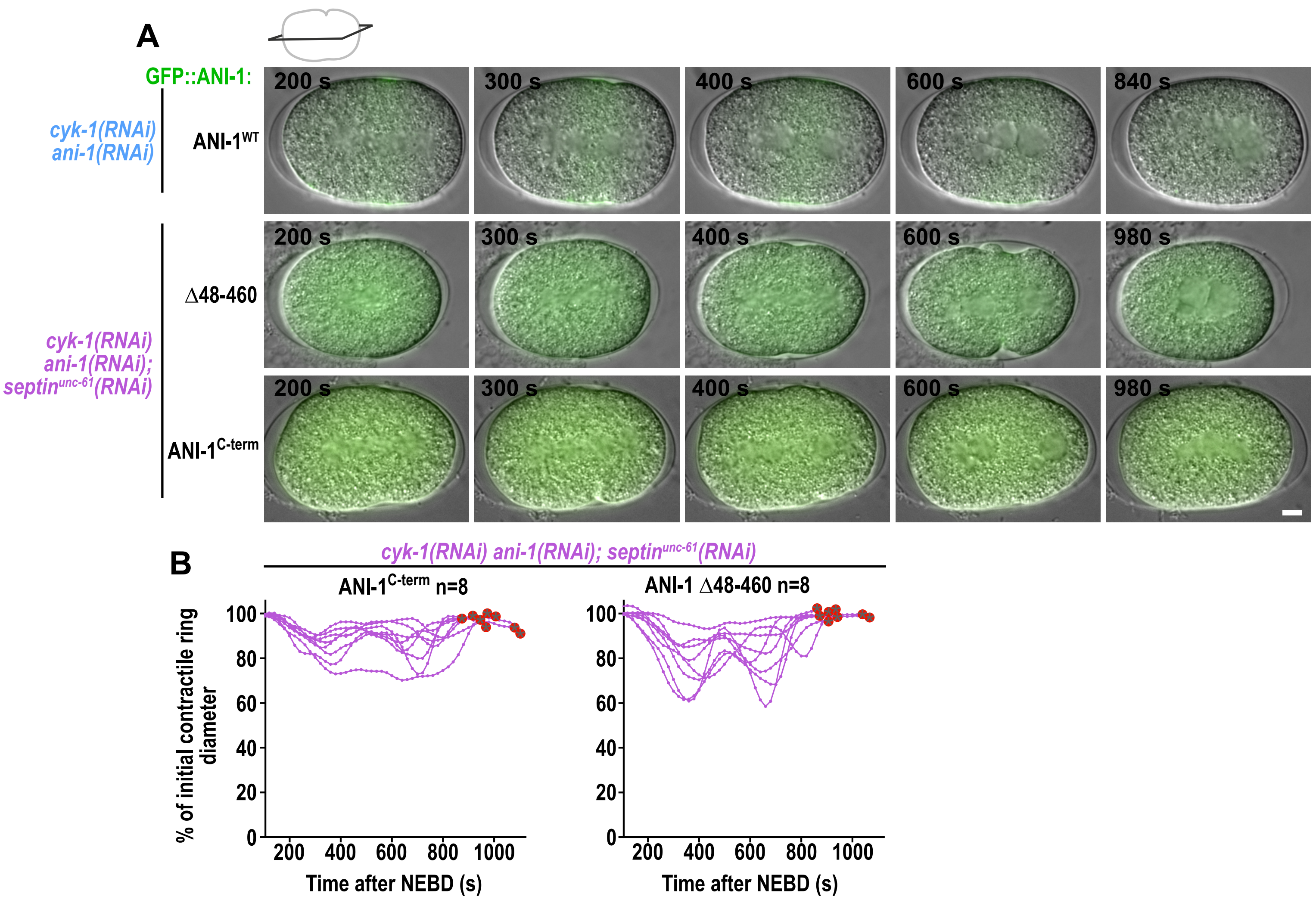

#### Supplemental FIGURE 11

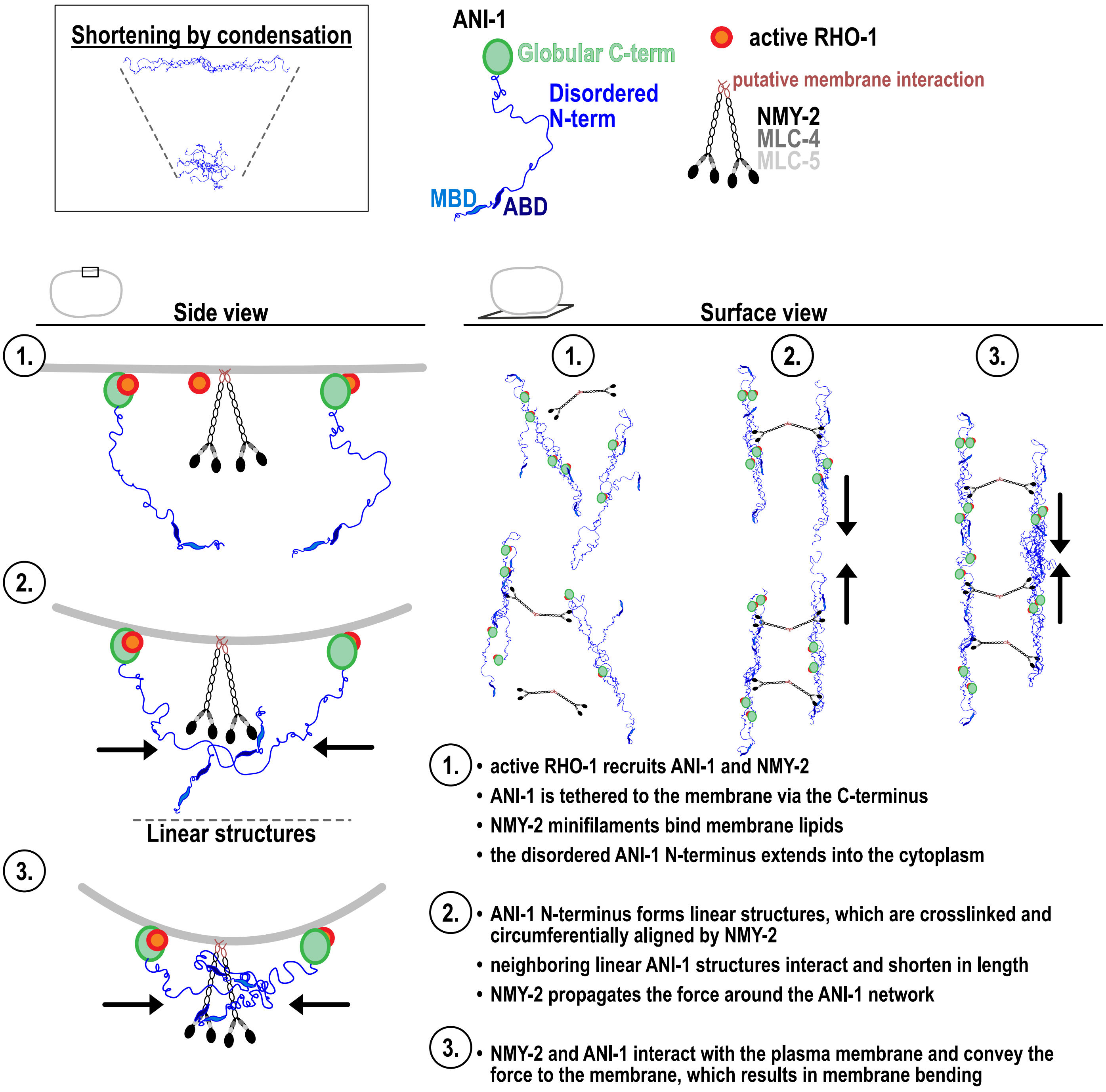
